# The impact of dietary fibres on gut microbiome of young children: insights from *ex vivo* experiments and an observational cohort

**DOI:** 10.64898/2026.02.11.703394

**Authors:** Shaillay Kumar Dogra, Norbert Sprenger, Dantong Wang

## Abstract

During weaning period, the intake of dietary fibres changes and increases dramatically. Given the considerable structural differences, we hypothesized that different fibres may vary in their function. The objective of the study was to explore the impact of specific dietary fibres (arabinoxylan, cellulose, pectin and xyloglucan) on the gut microbiome of children below 3 years. By using *ex vivo* fecal fermentation experiments, cellular models and cohort data analysis, we assessed how these fibres and their combinations influence infants’ gut microbiota composition, diversity, metabolite production and possible actions on the gut epithelial barrier function. We found that the fermentation with arabinoxylan, xyloglucan and pectin resulted in an increased production of short-chain fatty acids. These fibres also promoted the generation of metabolites with potential health benefits, such as indole-3-lactic acid. By combining the *ex vivo* fermentation and cellular co-culture experiments, arabinoxylan and xyloglucan were found to be able to maintain gut epithelial barrier integrity upon lipopolysaccharide challenge, and a blend of cellulose, pectin, and xyloglucan dampened different LPS induced cytokines. Moreover, pectin was found supporting the growth of a wide range of microbial species *ex vitro* and correlated positively with α-diversity in young children in an observational cohort. Our findings provided insights into the potential benefits of diverse fibre intakes during early life. Further studies are needed to understand the mechanisms and the effectiveness of specific fibres on the gut microbiome development in young children.

## INTRODUCTION

Infant feeding is critical for child development, and exclusive breastfeeding for the first 6 months of age is recommended by World Health Organization (WHO) to achieve optimal growth, development and health [1]. An important dietary transition is the introduction of complementary foods, which usually starts at around the age of 6 months followed by a gradual shift from first solid foods to family meals. During the process, a variety of foods are introduced to the children’s diet. The minimum diet diversity (MDD) has been used as one of the criteria to assess dietary quality for children aged 6-23 months, with grains, vegetables and fruits as key components of the recommended diet [1]. Consequently, fibres in children’s diet are also shifting from mainly human milk oligosaccharides (HMOs) to diverse types of dietary fibres [2].

Dietary fibres could be utilized by different gut bacteria depending on their carbohydrate-active enzyme (CAZyme) profile [3]. The selective fibre fermentation affects gut microbiome composition and function, influencing its development, important for gut health in early life [2]. One important aspect of the infant gut microbiome development is its diversification from 6 months onwards. Lack of age-appropriate gut microbiome diversification has been associated with increased non-communicable health risks, such as prediabetes and type II diabetes [4], atopic eczema [5], and vascular disorders [6]. In contrast, diverse gut ecosystems were found to be more resistant to environmental change [7], with lower risk for allergic diseases [8-10].

The main food sources of specific fibres are different. For example, arabinoxylan is a polysaccharide that can be found in various cereals, such as wheat, corn, and rice [11]. Cellulose is a component of plant cell walls, with high levels of presence in root and leafy vegetables, legumes, and some fruits, whereas pectin is commonly found in considerable amounts in fruits, especially in citrus fruits, pears and apples [12]. Xyloglucan is commonly present in the primary cell walls of higher plant [13], with *Tamarindus indica* seeds being a particularly rich source [14]. Due to the wide distribution of arabinoxylan, cellulose, pectin and xyloglucan in plant-based foods that are important in children’s diet, it is of interest to study their role in children’s microbiome development and potential health benefits. Although the role of dietary fibres on microbiome composition and function are increasingly studied in children, the exact impact of specific fibres is still unclear, as reviewed by Geniselli de Siva, *et al* and Noles, *et al* [15, 16].

Currently most studies in the field were conducted in adults. Since the microbiota composition changes rapidly in children under 3 years of age [2, 17], studies are needed to better understand the effect of specific fibres during gut microbiome development in early childhood and their potential health benefits. Additionally, to our knowledge there is very limited insight on the role of different fibres in the context of total diet, mainly due to the lack of dietary intake data for specific fibres.

In this study we aim to fill the knowledge gap by identifying relevant fibres that contribute to microbiome diversification in the postnatal age of 6 to 12 months and could have potential gut health promoting effects.

To do so, we combined *ex vivo* microbiome fermentation experiments with observational cohort analysis. First, we conducted an *ex vivo* experiment to systematically investigate the function of arabinoxylan, cellulose, pectin, and xyloglucan, both individually and in different combinations using fecal samples from 6 months and 12 months old donors and examined the formation of bene Short-Chain Fatty Acids (SCFA), beneficial metabolites, and barrier integrity and cytokine profile in a co-culture system after lipopolysaccharide (LPS) changes. Second, we analyzed dietary pectin intake in the total diet of normally developing young children under 3 years old in the Baby Connectome Project enriched (BCP-e) cohort [18], and investigated its association with microbiota α-diversity given its importance in child health [17].

## METHODS

### Test products

We previously identified four fibres constituting key fibres in early childhood diets that could influence microbiome development (data not shown). We sourced those as individual fibre products, containing over 95% cellulose, xyloglucan, arabinoxylan, or 80% pectin and tested them individually and in combinations. Inulin (IN, a prebiotic), arabinoxylan oligosaccharides (AXOS, a partially hydrolyzed arabinoxylan), and No Substrate Control (NSC, i.e., the fecal samples were incubated only with background medium) were included as controls.

Sources of these fibres were the following. Xyloglucan P-XYGLN and Arabinoxylan P-WAXYH were from Megazyme (Ireland); Inulin was from Sigma (United States). Cellulose Vitacel L600 was from J. Rettenmaier & Söhne GMBH(Germany); Pectin Herbacel SF 50-LV was from Herbafood (Germany); AXOS Soluble corn fibre SFC was from Agrifibre (United States).

Abbreviations used throughout the text: IN – inulin; A – AXOS; C – cellulose; X – xyloglucan; P - pectin, W – arabinoxylan; CX – combination of cellulose and xyloglucan with equal amount; CP – combination of cellulose and pectin with equal amount; XP – combination of xyloglucan and pectin with equal amount; CXP – combination of cellulose, xyloglucan and pectin with equal amount; XCPW – combination of xyloglucan, cellulose, pectin and arabinoxylan with equal amount.

### Ethical permission and donor selection

Fecal samples were collected according to a procedure approved by the Ethics Committee of the University Hospital Ghent (reference number BC-09977). Informed consent forms were signed by the parents of the children for donating fecal samples of their children.

Infants aged 6 or 12 months who were exclusively breastfed for 4 months and had already been introduced to complementary foods were eligible for study. Infants who had antibiotic use record within 90 days prior to fecal sample collection, history of disease such as Necrotizing enterocolitis (NEC) or gut surgery were excluded. In total 12 fecal samples were collected including 6 donors of 6 (±1) months and 6 donors of 12 (±1) months.

### *Ex vivo* experiment design

An *ex vivo* fecal fermentation study was performed using the SIFR^®^ technology (Cryptobiotix, Ghent, Belgium). The study was carried out as described previously [19]. In total, 12 fecal samples were incubated for 24 hours with 2 grams total fibre per liter of different fibres and fibre blends with equal amount for each fibre in the blends. Then the fundamental fermentation parameters, metabolites and microbiota composition were measured and spent supernatants prepared for subsequent testing on a Caco-2/THP-1 coculture system. A schematic overview of the experiment design is provided in Supplemental figure 1, Supplemental table 1 showed fibres and combination of fibres tested in the experiment.

### *Ex vivo* experiment measurements

#### Microbiota composition and fermentation parameters

Microbiota composition was measured by quantitative shallow metagenomics sequencing at 0h (only NSC) and 24h (all groups). Upon DNA extraction, standardized Illumina library preparation was performed followed by 3M total DNA sequencing reads. Results were analyzed at different taxonomic levels (species, family and phylum level). For taxonomic analysis, the proportional data derived from sequencing (%) was corrected for the total amount of cells present in each sample (quantified via flow cytometry)[19]. Community modulation score (CMS) was used to indicate changes at species level [20]. CMS represents the number of species (out of the 100 most abundant ones) that increased (positive CMS) or decreased (negative CMS) upon treatment. The combined CMS has a positive value when the number of increased species exceeds the number of decreased species, suggesting that the treatment is a diversity booster.

Fundamental fermentation parameters were measured, i.e., pH, gas, as well as SCFAs including acetate, propionate, butyrate, valerate, branched Chain Fatty Acids (bCFAs) and total SCFAs. Acetate, propionate, butyrate, valerate and bCFAs (sum of isobutyrate, isovalerate and isocaproate) were determined via a GC-FID approach [21].

#### Metabolomics

Liquid Chromatography-Mass Spectrometry (LC-MS) analysis was carried out using Thermo Scientific Vanquish LC coupled to Thermo Q Exactive HF MS. An electrospray ionization interface was used as an ionization source. Analysis was performed in negative and positive ionization mode. The Ultra Performance Liquid Chromatography (UPLC) was performed using a slightly modified version of the protocol described by Doneanu et al. [22]. Peak areas were extracted using Compound Discoverer 3.1 (Thermo Scientific). In addition to the automatic compound extraction by Compound Discoverer 3.1, a manual extraction of compounds included in an in-house library was performed using Skyline 21.1 (MacCoss Lab Software).

Identification of compounds were performed at three levels: Level 1-identification by retention times (compared against authentic standards), accurate mass (with an accepted deviation of 3ppm), and Tandem Mass Spectrometry (MS/MS) spectra; Level 2a-identification by retention times (compared against authentic standards), accurate mass (with an accepted deviation of 3ppm); Level 2b-identification by accurate mass (with an accepted deviation of 3ppm), and MS/MS spectra; Level 3 - identification by accurate mass alone (with an accepted deviation of 3ppm). Two hundred and seventy-eight metabolites were annotated on level 3, 46 on level 2b, 56 on level 2a, and 42 on level 1. Given the higher degree of certainty of correct annotation at level 1 and 2a, we mostly focused on level 1/2a metabolites.

#### Gut epithelial barrier and immune markers

Colonic epithelial Caco-2 cells obtained from the ATCC were cultured in Minimum Essential Medium (MEM) media supplemented with 10% Fetal Bovine Serum (FBS), 1X Non-Essential Amino Acids (NEAA) and 1mM Sodium Pyruvate. 24-well trans-well inserts were coated with Collagen I Rat Tail Protein and 1 x 10^5^ Caco-2 cells seeded onto the apical chambers. The basal chambers were filled with 500μl culture media. The plates were incubated at 37°C in a 5% CO_2_ humidified incubator for 14 days. During the differentiation process, culture media were changed every other day. Main experiments were performed when Trans-Epithelial Electrical Resistance (TEER) was more than 300 Ω.cm^2^.

THP-1 cells were cultured in RPMI-1640 supplemented with 10% FBS, 1mM sodium pyruvate and 10 mM HEPES at 37°C with 5% CO_2_. Cultures were initially inoculated at a density of 3 x 10^5^ cells/ml and split once density had reached 1 x 10^6^ cells/ml. To differentiate THP-1 cells into macrophages, THP-1 cells were centrifuged and re-suspended in cell culture medium containing 100 ng/ml Phorbol 12-myristate 13-acetate (PMA). The PMA-treated THP-1 cells were seeded (5 x 10^5^ cells) on transwell-suitable 24-well plates and incubated at 37°C in 5% CO_2_ to induce differentiation. After 48 hours, Caco-2 bearing inserts were moved to the transwell-suitable 24-well plates containing THP-1 cell derived macrophages (TPH-1m).

After appropriate differentiation periods for both Caco-2 and THP-1m, the co-culture experiment was performed in two steps: (i) a 24h treatment during which test products were applied on the apical side of the epithelial cells and (ii) a subsequent 6h lipopolysaccharide (LPS) challenge of THP-1m cells at the basal side to evaluate the impact of the test products on gut barrier integrity and immune cell response.

Colonic samples were centrifuged and filter-sterilized before use in the assay (administered apically as 20% of the cell medium: MEM medium supplemented with 1X NEAA and 1mM Sodium Pyruvate with 10% FBS, Gibco, Gibco, Carlsbad, CA, USA), then added to the apical chamber to replace the culture media. TEER was measured before and after LPS challenge. For the challenge, 500 ng/ml of LPS (Ultrapure LPS from E. coli 0111:B4, category number = tlrl-3pelps, Invitrogen, Carlsbad, CA, USA) was added to the basal chamber of the transwells containing the THP-1 cells and incubated for 6h. Samples from both apical and basal compartments were collected and subjected to cytokine/chemokine analysis using Multiplex Luminex^®^ Assay kit on the MAGPix^®^ analyser (for IL-6, CXCL10, IL-10, IL-1β, TNF-α) or ELISA (for IL-8).

### *Ex vivo* data analysis

Statistical analysis was conducted using a linear mixed-effects model, which accounted for both fixed effects (treatment) and random effects (interpersonal variation) (glmmTMB package v1.1.11 [23]. Effects of the treatments compared to the reference (NSC) were assessed via post hoc pairwise comparison using estimated marginal means (emmeans package v1.11.2). The Benjamini-Hochberg correction [24] was applied to control the false discovery rate (FDR) across parameters (stats package v3.6.2). For the statistical evaluation of the effects on microbial composition, the linear mixed-effects model was first applied to test overall effect of treatments on families/species, and only the families/species that displayed significant overall treatment effects were further considered for post hoc pairwise comparison. Effects were considered significant at p-adjusted ≤ 0.05. Statistical differences between treatments and NSC were indicated with ^*^ (0.01 < p-adjusted ≤ 0.05), ^**^ (0.001 < p-adjusted ≤ 0.01) or ^***^ (p-adjusted ≤ 0.001).

Consistently increased/decreased taxa were defined as any taxon that is present in at least four donors of a give donor group (either all, 6 months, of 12 months) that show a pattern of either increasing in all donors OR decreasing in all donors. The top 5 species were defined as those species that explain the most variation in a Principal Component Analysis (PCA) based on the log_2_ transformed absolute abundance matrix.

For statistical analysis of the quantitative shallow shotgun sequencing, a value below the limit of quantification (LOQ) was made equal to the LOQ. LOQ is one value per sample, determined by the read depth and total community density (LOQ of a sample = 1/total reads sequenced * density community). To be on the conservative side, we then removed all the data that is below the LOQ of the sample with highest LOQ [19]. The LOQ for us was 3.13 x 10^5^ cells/mL for 0h samples and 1.07 x 10^6^ cells/mL for 24h samples. Subsequently, statistics were performed based on log-transformation of the absolute values.

### Cohort data analysis

Dietary intake data and microbiota diversity of children aged 6 to 36 months from the BCP-e cohort were used in this study. BCP-e was a study conducted in the U.S. with a hybrid of accelerated longitudinal and cross-sectional study design which was described by Howell et al [18]. Research was approved by the Institutional Review Boards of University of North Carolina at Chapel Hill and University of Minnesota. Written informed consent was obtained from the parents. It included full term born infants with normal birth weight and absence of major pregnancy and delivery complications. Energy and nutrient intakes, including pectin intake, were estimated based on 2 days 24h recall, only data collected within 3 days prior to fecal sample collection were included in the analysis. Subjects with energy intake exceeded 3 standard deviations (SD) were removed. Shannon diversity index was used as a microbiota α-diversity measure. A linear mixed effect model was applied to analyze the association between pectin intake and Shannon diversity index. Subject numbers were used as random effects. Several models were tested with different covariate adjustments, including age, energy, and total as well as soluble fibre intakes.

## RESULTS

### Treatment with arabinoxylan and xyloglucan increased the abundance of adult-type species whereas treatment with pectin increased community modulation score

The change of pH and gas formation that serve as proxy for microbial fermentation were monitored first. Compared to NSCs and baselines, all tested fibres, including inulin and AXOS, decreased the pH after 24h incubation, albeit slightly less with cellulose, and increased gas production (Supplemental figure 2).

Next, we investigated the effect of specific fibres on microbiota composition. After a 24h incubation, both arabinoxylan and xyloglucan stimulated adult-type *Bifidobacterium* species, such as *B. catenulatum, and B. pseudocatenulatum* in 12 months donor samples, as illustrated by changes from baseline (Supplemental figure 3 and Supplemental data file 1). On the contrary, the infant-type *Bifidobacterium* species, such as *B. longum, B. bifidum* and *B. breve*, were not increased upon any of the fibre treatments, except for *B. longum* upon incubation with arabinoxylan (Supplemental figure 3 and Supplemental data file 1).

Pectin or the combination of pectin and xyloglucan significantly increased the abundance of 16 species mainly at 12 months, with 10 out of 16 being common between the two treatments, which was the highest number of species that were significantly increased among the evaluated fibres (Supplemental figure 3 & Supplemental data file 1). For Pectin, some of the increased species were *Bifidobacterium longum, Bifidobacterium catenulatum, Bifidobacterium pseudocatenulatum, Faecalibacterium prausnitzii, Phocaeicola dorei, Veillonella ratti* and *Bacteroides ovatus*. The highest significant fold-change was observed for *Faecalibacterium prausnitzii* and then for *Phocaeicola dorei* at 12 months. The combination of xyloglucan and pectin also significantly increased 16 different species, including *Bifidobacterium catenulatum, Bifidobacterium longum, Bifidobacterium pseudocatenulatum, Blautia wexlerae, Faecalibacterium prausnitzii, Phocaeicola vulgatus, Veillonella dispar*, and *Veillonella parvula*. No significant change was found for cellulose (Supplemental figure 3 & Supplemental Data file 1).

As a proxy for diversity, we calculated the Community modulation score (CMS) upon incubation with the different fibres and their blends. As shown in figure 1, compared to NSC, no statistically significant difference was found at 6 months except for the combination of cellulose and pectin. At 12 months, both inulin and AXOS promoted the growth of multiple species. Among tested fibres, arabinoxylan and xyloglucan supported the growth of fewer species, whereas treatment with pectin and the combination of pectin with cellulose or xyloglucan resulted in the growth of the greater number of species, when incubated with samples from 12 months children (Figure 1).

**Figure 1.**
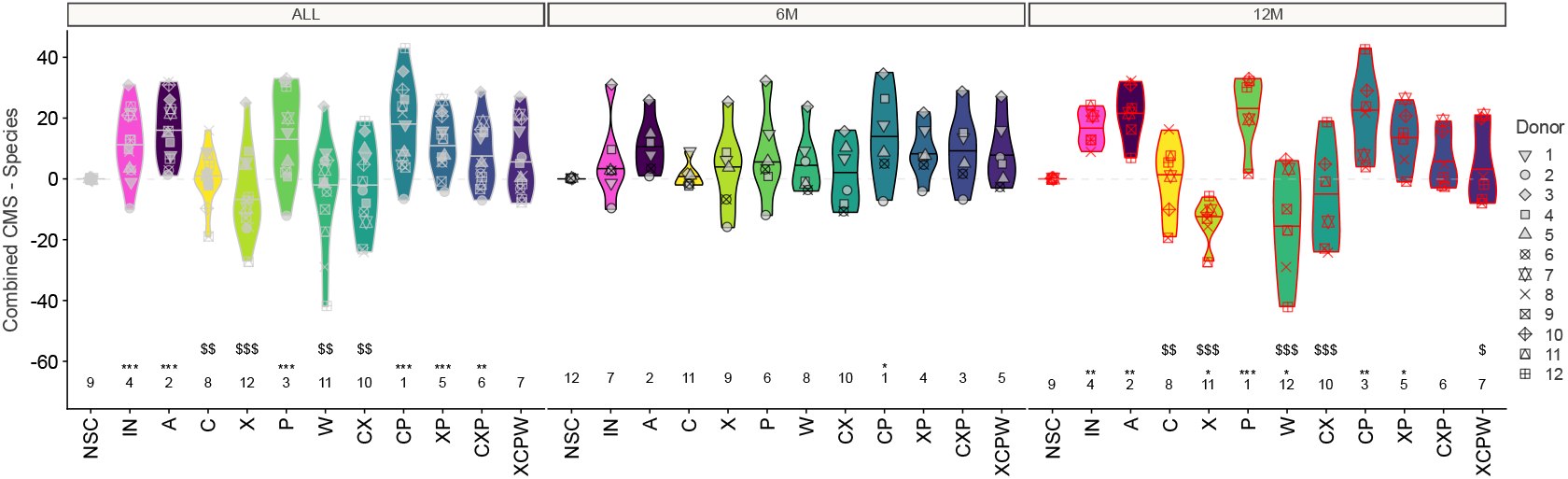
The impact on the combined community modulation score (CMS) at species level. Samples were collected after 24h of incubation. Statistical differences between NSC and the individual treatments are visualized via ^*^ (0.01 < p-adjusted < 0.05), ^**^ (0.001 < p-adjusted < 0.01) or ^***^ (p-adjusted < 0.001). Significant differences between Inulin and individual treatments are indicated via $/$$/$$$. The rank of the average values per treatment are indicated at the bottom of the figure. Ranks comprise here the median of the log-fold changes, not the raw data.

### Fermentation of arabinoxylan, pectin and xyloglucan increased beneficial gut metabolites

As shown in Figure 2, inulin treatment significantly increased total SCFAs, particularly acetate and propionate levels (Supplemental figure 4), whereas cellulose had no significant effect on total SCFAs as compared to NSC. The treatments with pectin, arabinoxylan and xyloglucan, as well as their combinations with cellulose, significantly increased the production of total SCFAs, mainly driven by the increase of acetate and propionate, especially at 12 months. No significant increase in butyrate was observed (Supplemental figure 4).

**Figure 2.**
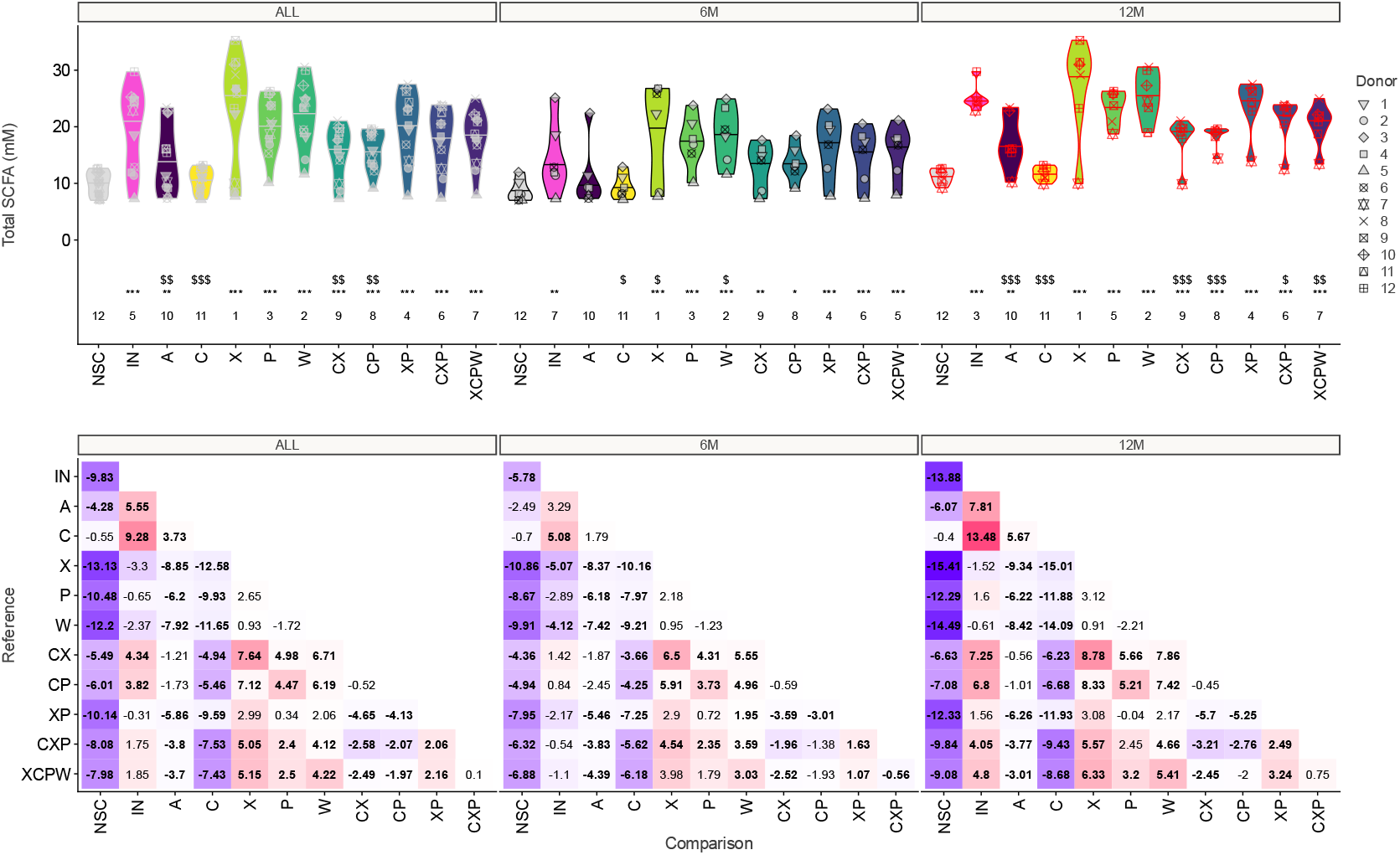
Impact of fibres on SCFAs production. Top panel: The impact of the tested products on total SCFA for 12 infants (6 months old (n= 6); 12 months old (n = 6)). Statistical differences between NSC and the individual treatments are visualised via * (0.01 < p-adjusted < 0.05), ** (0.001 < p-adjusted < 0.01) or *** (p-adjusted < 0.001). Significant differences between inulin and the individual treatments are indicated via $/$$/$$$. The rank of the average values per treatment are indicated at the bottom of the figure. Bottom panel: Values in each matrix represents the difference (in mM) between the product on the horizontal axis compared to the one on the vertical axis, with significant differences (p-adjusted < 0.05) being indicated in bold.

Compared to NSCs, treatment with AXOS, pectin and arabinoxylan showed significant higher indole-3-lactic acid (ILA) concentrations in samples from 12 months infants, while a slight increase was found by the combination of pectin with cellulose, as well as by the combination of the 4 fibres, pectin, cellulose, xyloglucan and arabinoxylan (Figure 3). Similarly, Indole-3-methyl acetate significantly increased in samples from 12 months infants, but not in 6 monthsinfants, upon various treatments, particularly for fibres arabinoxylan and xyloglucan (Supplemental figure 5 and Supplemental data file 2).

**Figure 3.**
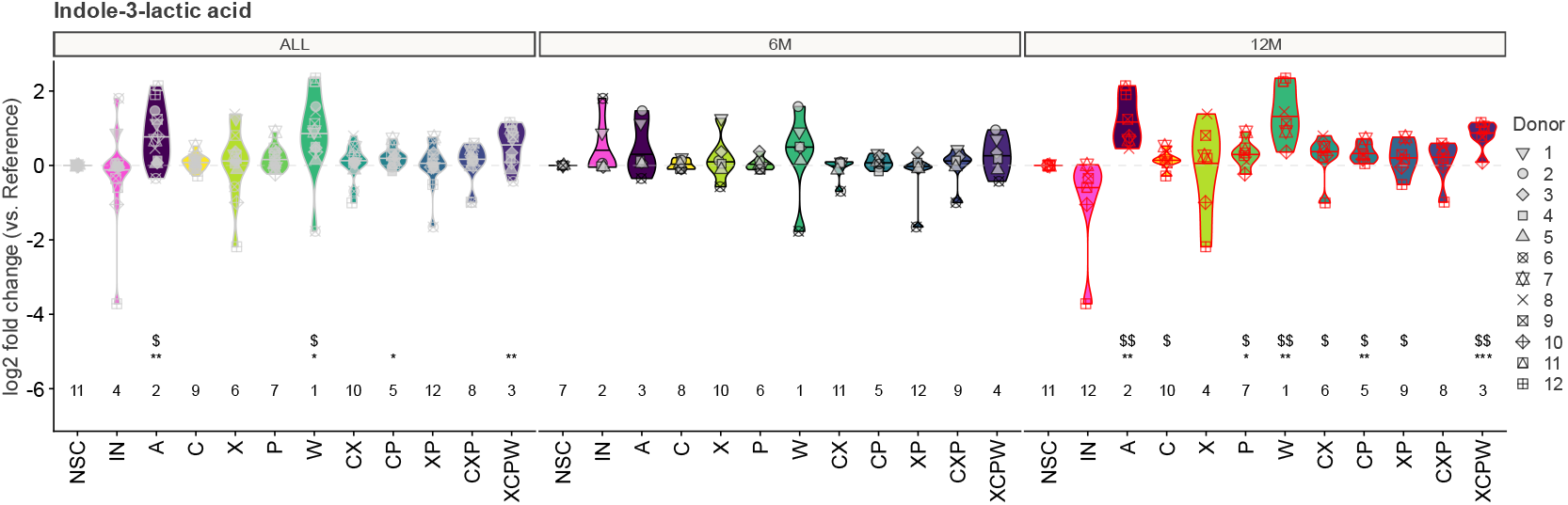
The impact of the test products on indole-3-lactic acid for 12 infants (6 months old (n= 6); 12 months old (n = 6)). Statistical differences between NSC and the individual treatments are visualised via * (0.01 < p-adjusted < 0.05), ** (0.001 < p-adjusted < 0.01) or *** (p-adjusted < 0.001). Significant differences between IN and the individual treatments are indicated via $/$$/$$$. The rank of the average Log2FC per treatment are indicated at the bottom of the figure. Ranks comprise here the median of the log-fold changes, not the raw data.

Acetylated amino acids significantly increased upon fibre fermentation, particularly N-acetylated amino acids in samples from 12 months infants (Supplemental figure 6). Inulin, xyloglucan and the combination of xyloglucan and cellulose most strongly enhanced the production of N-acetylaspartic acid (Supplemental figure 6A); N2-acetyllysine was significantly increased especially at 12 months by every treatment except cellulose (Supplemental figure 6B); and N-acetyl histidine significantly increased strongest under arabinoxylan and inulin and slightly under AXOS fermentation, as well as with the combinations of cellulose, xyloglucan and pectin or cellulose, xyloglucan, pectin, and arabinoxylan (Supplemental figure 6C).

At 6 months only, 2-hydroxyisocaproic acid (HICA) significantly increased upon the fermentation of inulin, xyloglucan, arabinoxylan and the combinations of cellulose and xyloglucan; xyloglucan and pectin; cellulose, xyloglucan and pectin; xyloglucan, cellulose, pectin and arabinoxylan (supplemental figure 7A). At 12 months, the production of pipecolinic acid (PIPA) was increased by all tested fibres and fibre mixes to different extents (Supplemental figure 7B). At 12 months, acetyl-agmatine significantly increased upon all tested fibres and fibres mixes, except cellulose, while at 6 months it significantly increased with pectin, arabinoxylan, and combinations of cellulose and xyloglucan; xyloglucan and pectin; cellulose, xyloglucan and pectin; xyloglucan, cellulose, pectin and arabinoxylan (Supplemental figure 7C). Similarly to inulin at 12 months, compared to the NSC, trimethylamine N-oxide (TMAO) significantly increased upon all tested fibres and fibre mixes, except cellulose, while only the combination of xyloglucan and pectin at 6 months showed a significant increase (Supplemental figure 7D). Kynurenic acid (KYNA) increased significantly after treatment with various fibre and fibre mixes in samples from 12 months infants, except for cellulose plus xyloglucan, and mix of xyloglucan, cellulose, pectin and arabinoxylan (Supplemental figure 7E).

Regarding vitamins, nicotinamide (a form of Vitamin B3), slightly but significantly increased at 12 months under treatments with inulin, pectin, arabinoxylan and for the combinations of cellulose and pectin; cellulose, xyloglucan and pectin; xyloglucan, cellulose, pectin and arabinoxylan (Supplementary figure 5). Also, pyridoxamine (vitamin B6) increased significantly by pectin and the combination of cellulose and pectin for 12 months infants. Trigonelline, a product of nicotinic acid (vitamin B3) metabolism, tended to increase at 12 months, particularly after the treatments of AXOS, cellulose, pectin, arabinoxylan, and combinations of cellulose and xyloglucan; cellulose and pectin; cellulose, xyloglucan and pectin (Supplemental figure 5).

### Arabinoxylan treatments increased trans-epithelial electrical resistance (TEER) upon LPS challenge, and altered cytokine profiles

A Caco-2/THP-2 co-culture system was used to test the potential function of spent culture media after fibre fermentation on gut epithelial barrier function. As shown in Figure 4, all individual fibres, except inulin and cellulose, tended to increase TEER at 12 months as compared to NSC, while none reached statistical significance. For 6 months infants, xyloglucan and arabinoxylan exhibited stronger effects with the latter being significant (Figure 4).

**Figure 4.**
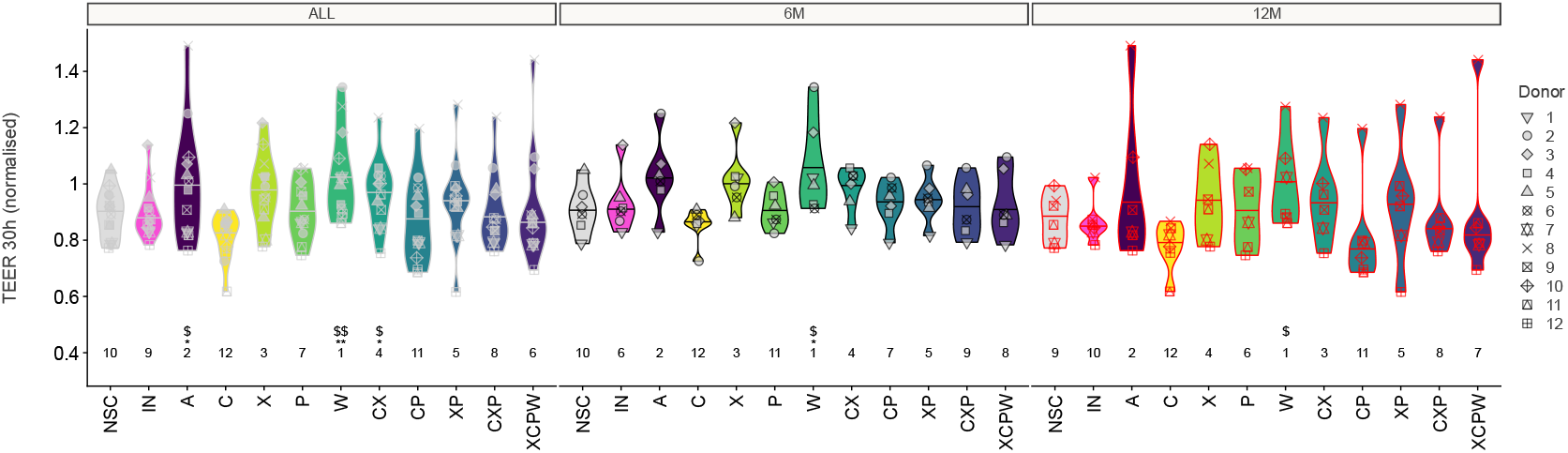
The impact of the test products compared to a no substrate control (NSC) on gut barrier integrity as measured by the TEER of the Caco-2 epithelial layer and 30h treatment (24-hour incubation without LPS followed by 6-hour treatment with LPS) in the basal compartment containing differentiated THP-1 cells) using the SIFR^®^ technology. Samples exposed to the cells were collected from the SIFR^®^ model after 24h of colonic incubation. Statistical differences between NSC and the individual treatments are visualised via * (0.01 < p-adjusted < 0.05), ** (0.001 < p-adjusted < 0.01) or *** (p-adjusted < 0.001). Significant differences between inulin and the individual treatments are indicated via $/$$/$$$. The rank of the average value per treatment are indicated at the bottom of the figure.

A correlation analysis was performed between the individual SCFA after 24h fibre incubation and TEER. Acetate, propionate and total SCFA were found correlated with the increase of TEER, especially at 12 months (Supplemental figure 8). A positive correlation was also observed between increased TEER and the presence of, among a few others, acetate/lactate-producers, such as *Bifidobacterium_u_s* and, *Bifidobacterium catenulatum*, which were also increased after the treatment of arabinoxylan, pectin and xyloglucan (Supplemental figure 9).

For 6 months infants, the combination of cellulose, xyloglucan, and pectin significantly exhibited lower levels of both anti-inflammatory (such as IL-10) and pro-inflammatory cytokines (such as IL-6, TNF-α, IL1-β and CXCL-10) (Supplemental figure 10). Additionally, TNF-α for 6 months infants was significantly lowered by cellulose, xyloglucan, and the combination of xyloglucan and pectin when compared to the NSCs. At 12 months, only cellulose significantly increased IL-10 and CXCL-10 concentrations (Supplemental figure 10).

### Consumption of pectin associated with gut microbiota α diversity in young children

The BCP-e cohort of normally developing children captured food intake data allowing to deduce total-and soluble fibre intake as well as pectin intake per day per child. Unfortunately, the quantitative data coverage for other fibre types was below 45% among fibre containing foods (data not shown). Since we observed that pectin fermentation promoted the growth of the largest number of species among tested fibres, we analyzed the relationship between pectin intake and microbiota α-diversity using data from BCP-e cohort. In total 256 subjects were included in the cohort data analysis. A positive association was found between the amount of pectin intake (g/day) and Shannon index, adjusted for age in months, energy and soluble fibre intakes (Table 1). A similar positive association was also obtained for the proportion of pectin in the total soluble fibre intake (coefficient=0.03, p<0.001, not shown). Other diversity index was also tested and the same trend was found (Supplemental table 2).

**Table 1.** The association between pectin intake and microbiota α-diversity in cohort study.

|  | Shannon |  |  |  | ShannonH |  |  |  |
| --- | --- | --- | --- | --- | --- | --- | --- | --- |
|  | Coefficient | 95% CI |  | P | Coefficient | 95% CI |  | P |
|  |  | Lower | Upper |  |  | Lower | Upper |  |
| Pectin (g/d) |  |  |  |  |  |  |  |  |
| Model 1 | 0.103 | 0.067 | 0.139 | <0.001 | -0.056 | -0.175 | 0.062 | 0.352 |
| Pectin (g/d) |  |  |  |  |  |  |  |  |
| Model 2 | 0.043 | 0.005 | 0.082 | 0.028 | -0.001 | -0.132 | 0.13 | 0.987 |
| Pectin (g/d) |  |  |  |  |  |  |  |  |
| Model 3 | 0.073 | 0.021 | 0.126 | 0.007 | -0.009 | -0.188 | 0.17 | 0.924 |
| % pectin in<br>soluble fiber |  |  |  |  |  |  |  |  |
| Model 4 | 0.031 | 0.015 | 0.047 | <0.001 | 0.007 | -0.048 | 0.062 | 0.803 |

Linear mixed effect models were used for the analysis. Subject number was used as random effect.
Model 1 adjusted for age; Model 2 adjusted for age and energy intake; Model 3 adjusted for age, energy and soluble fibre intakes; Model 4 each 10% of pectin in total soluble fibre adjusted for age, energy and soluble fibre intakes. Shannon: Shannon index. ShannonH: Shannon index after removal of bacteria modulated by pectin found in *ex vitro* experiment.

To test the robustness of the observed associations, we recalculated the Shannon index (ShannonH) without the taxa that we observed growing upon pectin treatment in the preclinical model (i.e. Bacteroides (*Bacteroidaceae, Bacteroidales_u_f, Bacteroides thetaiotaomicron, Eubacterium eligens, and Faecalibacterium prausnitzii*). After removing pectin promoted taxa we lost the significant association with pectin, suggesting the association between pectin intake and microbiota α-diversity is likely mediated by those microbial species. It is worth noting that we also analyzed the relationship between total soluble fibre intake and microbiota α-diversity, no significant result was obtained (data not shown).

## DISCUSSION

The weaning period is a critical transition phase during child development. Studies have shown that the introduction of a variety of foods can help not only to shape the child’s palate and encourage long lasting healthy eating patterns [25], but also to establish a diverse and healthy microbiome, which plays an important role in the overall health and immune system of children [7, 26]. The current WHO complementary feeding guidelines highlights the importance of food diversity, and fibre rich foods, such as grains, fruits and vegetables are among the 8 food groups for diet diversity [1]. Due to the differences in molecular structure, the functions of different fibres are different. In this study, we investigated the impact of specific fibres (cellulose, xyloglucan, pectin, and arabinoxylan), individually and in combinations, on gut microbiome composition and function using samples from 6 months and 12 months old donors, aiming to provide insights on their specific effects on the gut microbiome including its functional potential, which is important for long term health [2].

The functions of pectin have been investigated in several studies. For example, it has been reported that human gut microbiota composition and metabolites modulated by a combination of citric pectin and *Bifidobacterium longum BB-46 in vitro* which implied the potential prebiotic function of pectin [27], but another study showed that sugar beet pectin supplementation did not alter profiles of fecal microbiota in healthy adults [28]. The inconsistent results suggested that the influence of pectin may depend on its specific food source which reflecting the difference in the structure. In a systematic review, Pascale *et al* summarized multiple lines of evidence and demonstrated that pectin fermentation *in vitro* could increase the production of SCFAs, especially acetate, which is consistent with our findings [29]. The microbiota composition also changed, either directly by pectin fermentation or by cross-feeding interactions, and resulted in the increased abundances of multiple bacterial families, such as *Ruminococcaceae, Bacteroides* and *Lachnospira genera* and species such as *Bifidobacterium spp*. and *Faecalibacterium prausnitzii* a.[29]. In our study, pectin significantly increased the abundance of 16 species and particularly boosted the SCFAs producers, such as acetate/propionate producing *Phocaeicola dorei* and *Phocaeicola vulgatus*, and butyrate-producing *Flavonifractor plautii* and *Faecalibacterium prausnitzii* which is consistent with a previous study [29].

For the effect of fibres on gut microbiota diversity, the *in vitro* study results were not consistent either. Studies suggested that the microbiota Chao1 (species richness) and Shannon index (α-diversity) was either no change or decreased as expected if individual taxa grows higher[27, 28]. This might be because the *in vitro* experiments were usually conducted in a closed environment, no new species were introduced. To overcome this issue, we used the number of growing species to indicate the potential effect of each fibre on diversifying microbiota. Pectin was found to promote the growth of the highest number of species among tested fibres suggesting its potential in promoting microbiota diversity *in vivo*. This finding was strengthened by an observational cohort data analysis in which a positive association between pectin intake and α-diversity was observed. When total soluble fibre was analyzed, no significant outcome was obtained. In addition, the association between pectin and microbiota α-diversity disappeared when the bacterium species influenced by pectin in *ex vivo* fermentation experiment were removed from the analysis. This feature renders a certain degree of specificity to pectin. Diverse microbiotas are particularly beneficial during early childhood as they contribute to better health outcomes [7, 26]. To our knowledge, this is the first report on the potential effect of pectin in promoting microbiota diversity in young children. It underscores the importance of pectin consumption in the weaning period.

Based on high abundance in children’s diets and sparse knowledge of their microbiome modulating effects, we also studied arabinoxylan, cellulose and xyloglucan. We found that arabinoxylan and xyloglucan treatment increased SCFAs, N-acetylated amino acids and indole-3-lactic acid, especially at 12 months. An improved barrier integrity was also observed after fermentation with arabinoxylan, and to a lesser extent with xyloglucan, after a 6h LPS challenge. On the other hand, xyloglucan showed a more pronounced effect on released cytokines. These effects might be attributed to the specific metabolite profiles obtained after fermentation by children’s gut microbiota. SCFAs have been shown to be important for maintaining gut health by providing energy to colonocytes, reducing inflammation, and enhancing the integrity of the gut barrier [30]. Specifically, acetate and propionate have been reported to have a protective role against respiratory problems, butyrate and propionate play a role in intestinal homeostasis [31]. Amino acid-derived metabolites, like indole-3 lactic acid and others play a wide range of roles in both the human body and the gut microbiome, influencing immune responses and other physiological processes [25]. Indole-3-lactic acid mediates host-microbial crosstalk for immune response [32] and was reported for its protection against gut infections [33-35]. Its anti-inflammation function has been showed *in vitro* and fecal Indole-3-lactic acid level was found negatively correlated with the progression indicator of inflammatory bowel diseases [36]. These reported results together with our findings suggest that arabinoxylan and xyloglucan modulate children’s microbiome in a beneficial manner.

In this study, compared to NSCs, we found TMAO increased with all tested fibres, except cellulose. TMAO was associated with childhood obesity [37] and stroke risk in adults [38]. It is a typical metabolite derived from animal-based foods, such as red meat consumption. However, inconsistent results have been reported for the relationship between dietary fibres and TAMO levels. Some studies reported that fibre rich plant-based diet intake could counterbalance TMAO formation [39, 40], but Almer *et al* reported that TMAO concentrations naturally increased from childhood to early adults, and fibre intake was positively associated with serum TMAO in females but not in males [41]. Further research is needed to investigate the role of fibre in TMAO metabolism.

Different dietary fibres co-exist in the diet and influence fermentation, microbiome modulation and other related functions. Therefore, one important aspect in investigating the function of dietary fibres on microbiome development and gut health is to evaluate the function of fibre combinations. To investigate the effect of specific fibres in foods where multiple fibres coexist, we tested several fibre combinations. A fibre blend containing cellulose, pectin, and xyloglucan was shown to lower both pro-and anti-inflammatory cytokine levels, thus deescalate overall immune reaction. To our knowledge this is the first study reporting this effect. Given the beneficial effect of these fibres on SCFAs, microbiota diversity and barrier integrity, we hypothesize that the overall effect would be beneficial, but the exact implication is yet to be investigated.

This study is not without limitation. First, the results obtained in this study were derived from the use of pure fibres rather than foods or ingredients that are naturally rich in these fibres. This distinction is important because the effects observed with isolated fibres may not fully replicate the complex interactions that occur when these fibres are consumed as part of whole foods. Secondly, the fibres were tested in an *in vitro* experimental system with a small sample size, which may limit the generalizability of the findings. Furthermore, the fecal samples collected at 6 and 12 months were from different donors, which might introduce higher inter person variability and affect the consistency of the results.

## CONCLUSION

In this report we highlighted that pectin enhanced SCFAs production and modulated gut microbiota composition. It promoted the growth of multiple species in *ex vivo* experiments and was positively associated with microbiome α-diversity in an observational cohort study. Arabinoxylan and xyloglucan also contributed to SCFAs production and improved gut barrier integrity. A fibre blend of cellulose, pectin, and xyloglucan modulated cytokine profile after LPS challenge. These findings provided insights on specific functions of each tested fibre. Further research is needed to understand the mechanisms and actual health benefits.

## ACKNOWLEDGEMENTS

The authors thank Dr. Jian Zhang for his contribution in the BCP-E cohort data analysis, and Professor Weili Lin for his guidance on the BCP-E data structure and result interpretation. We acknowledge Drs Jovyn Frost and Ana Rovalino for sharing their expertise in dietary fibre research, and Drs. Giles Major and Marie Bachelet for organizing the study, Jonas Poppe for analyzing *ex vivo* experiment data, and Pieter van den Abbeele for his contribution to related scientific discussions.

## Conflict of interest statement

No conflict of interest. SKD, NS and DW are current employees of Société des Produits Nestlé SA (Vevey, Switzerland).

## Notes

### Competing Interest Statement

All authors are employees of Nestle Research, Societe des Produits Nestle SA.

### Funding

- Nestle Research

