## Supplemental Tables 1-2, Supplemental Figures 1-10 for "The impact of dietary fibres on gut microbiome of young children: insights from *ex vivo* experiments and an observational cohort"

### SUPPLEMENTARY INFORMATION

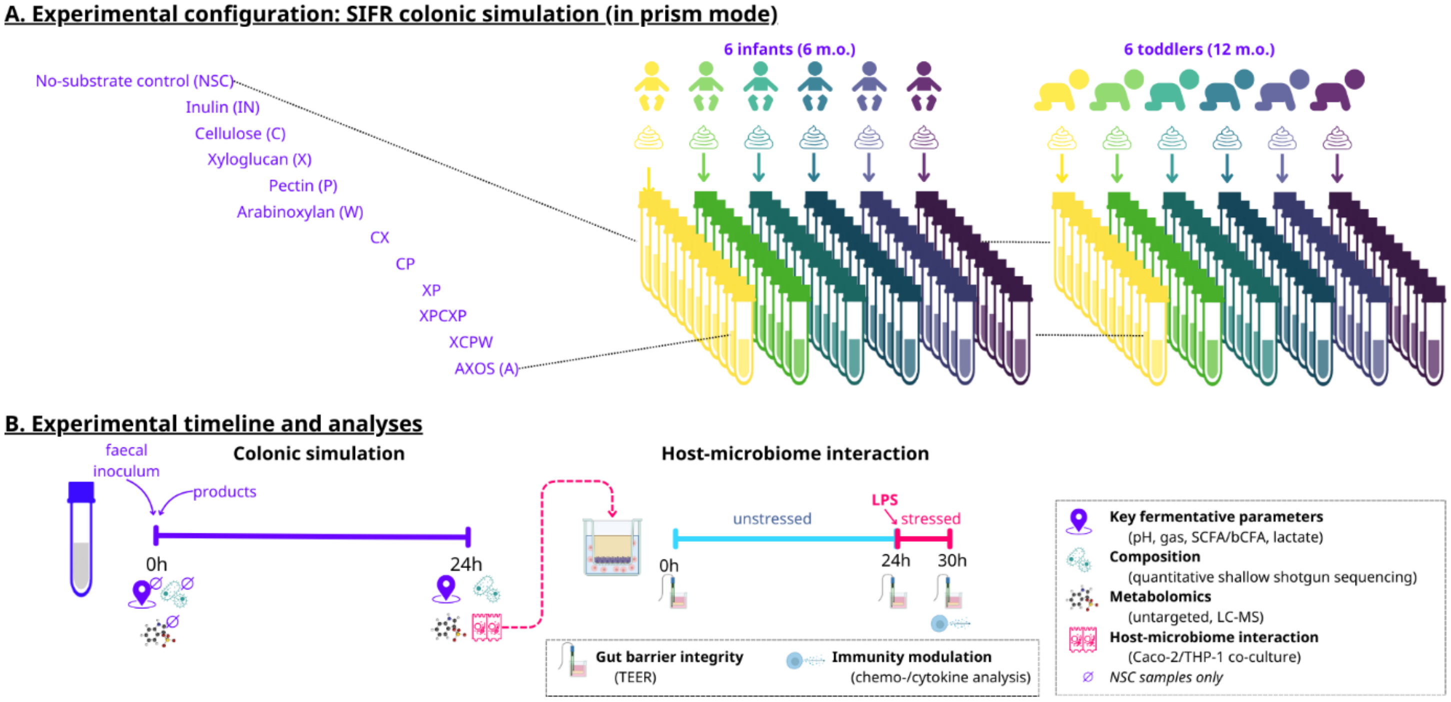


**Supplemental Figure 1** - Schematic overview of the *ex vivo* study design, timeline and analyses. SIFR® technology platform was used to assess the impact on the gut microbial activity and composition of 12 infants (6 months old (n= 6); 12 months old (n = 6)) of specific types of fibres and their combinations (C, X, P, W, CX, CP, XP, CXP, CXPW, A) compared to a no substrate control (NSC) and a reference prebiotic (IN). Panel A. Experiment design. Panel B. Timeline and analyses implemented in the *ex vivo* study. Abbreviations: IN - Inulin, A - AXOS, C - Cellulose, X - Xyloglucan, P - Pectin, W – Arabinoxylan, CX – Cellulose & Xyloglucan, CP – Cellulose & Pectin, XP – Xyloglucan & Pectin, CXP – Cellulose & Xyloglucan & Pectin, XCPW – Xyloglucan & Cellulose & Pectin & Arabinoxylan.

**Supplemental table 1. The matrix of tested fibers and their combinations.**

| Tested fibers | Cellulose (C) | Pectin (P) | Xyloglucan (X) | Arabinoxylan (W) |
|---|---|---|---|---|
| Cellulose (C) | 2 g/L | | | |
| Pectin (P) | | 2 g/L | | |
| Xyloglucan (X) | | | 2 g/L | |
| Arabinoxylan (W) | | | | 2 g/L |
| CP | 1 g/L | 1 g/L | | |
| CX | 1 g/L | | 1 g/L | |
| XP | | 1 g/L | 1 g/L | |
| CXP | 0.66 g/L | 0.66 g/L | 0.66 g/L | |
| CXPW | 0.5 g/L | 0.5 g/L | 0.5 g/L | 0.5 g/L |

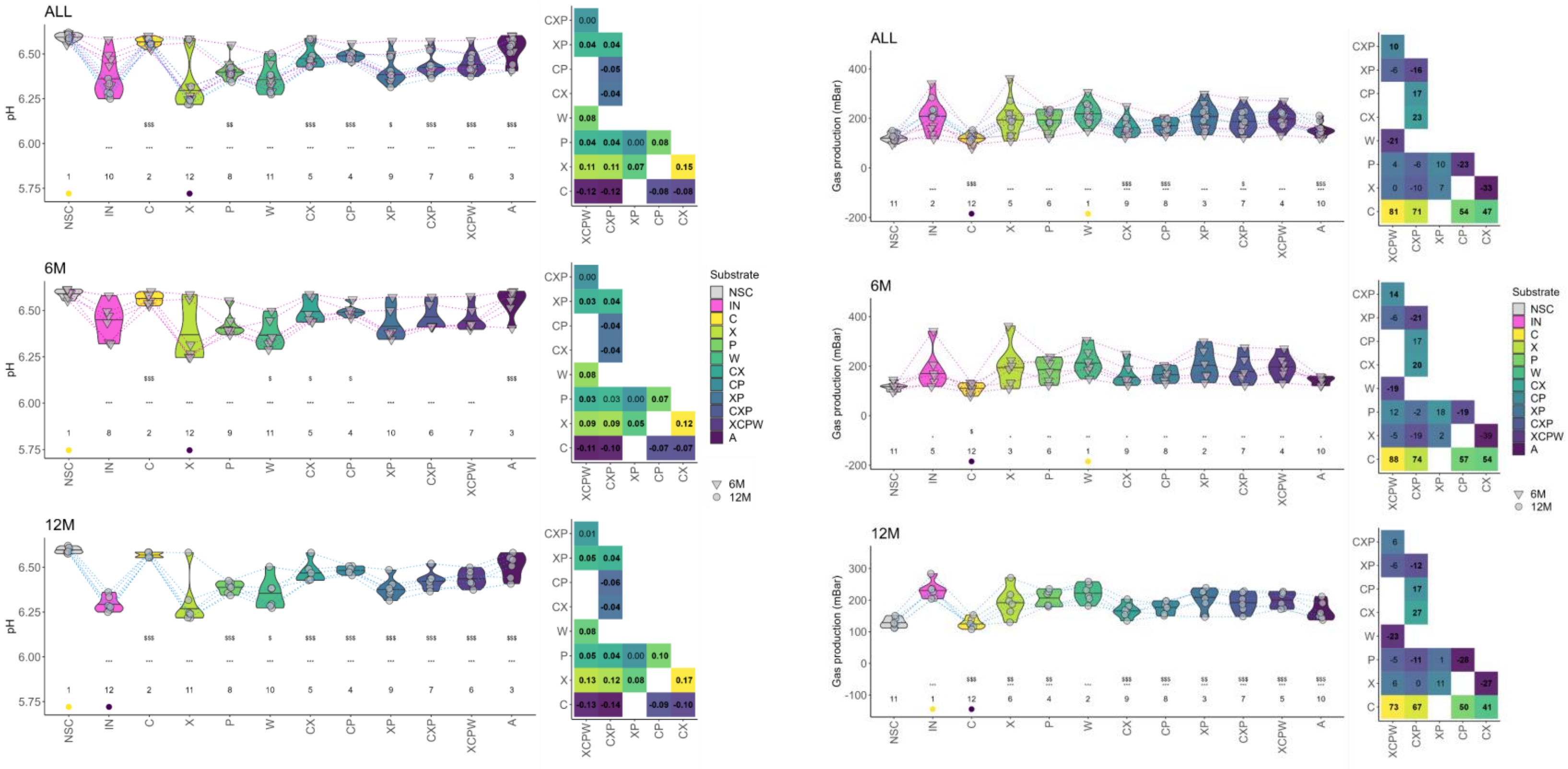


**Supplemental Figure 2** – Measurements of pH and Gas production in the *ex vivo* experiment. Abbreviations: IN - Inulin, A - AXOS, C - Cellulose, X - Xyloglucan, P - Pectin, W – Arabinoxylan, CX – Cellulose & Xyloglucan, CP – Cellulose & Pectin, XP – Xyloglucan & Pectin, CXP – Cellulose & Xyloglucan & Pectin, XCPW – Xyloglucan & Cellulose & Pectin & Arabinoxylan.

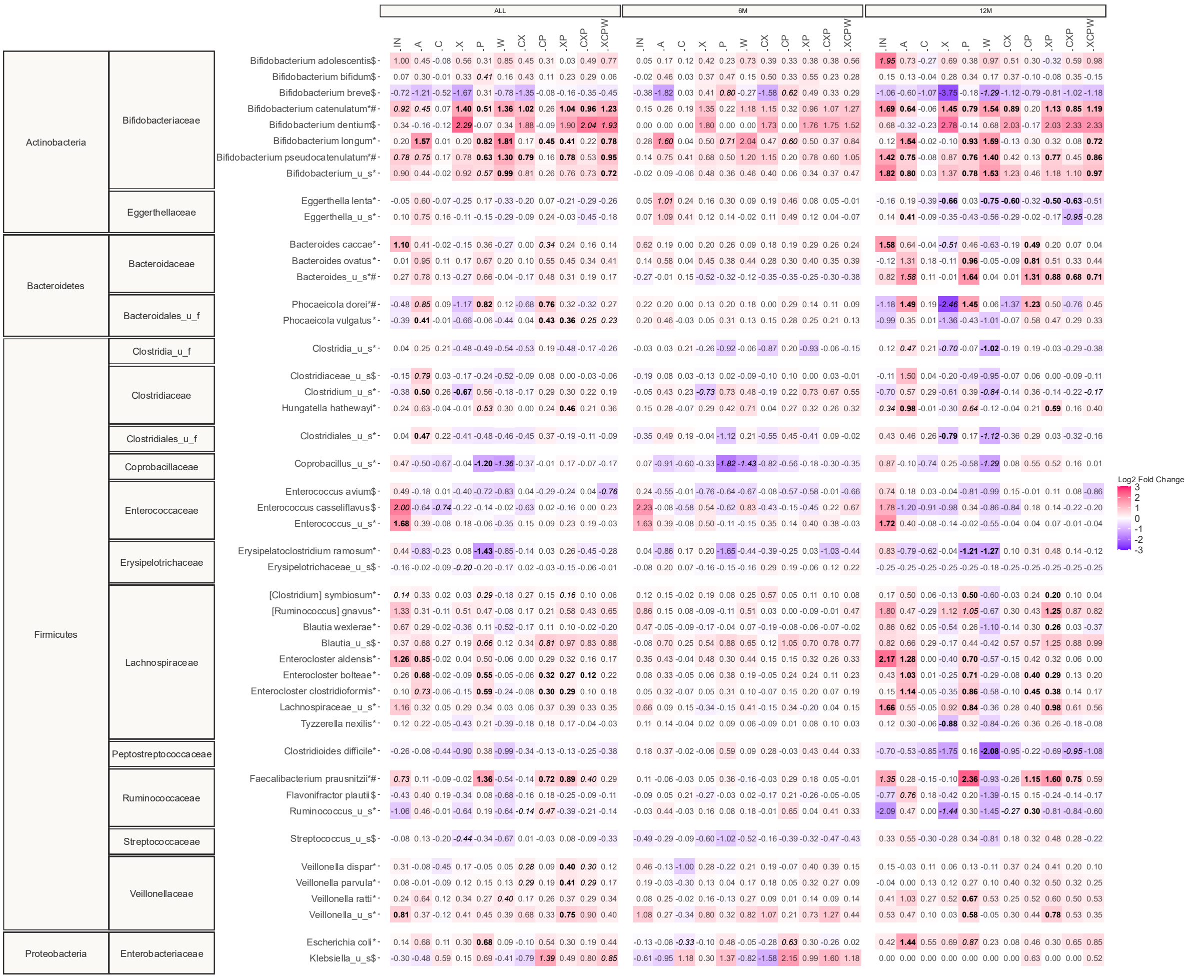


**Supplemental Figure 3.** The impact of the test products on bacterial species that were significantly affected (padjusted < 0.05, indicated with *) or consistently affected for at least 4 donors (indicated with $) by any of the treatments, expressed as $log_2$ transformation of abundance treatment/abundance NSC ratios ($log_2FC$), averaged across all donors. Samples were collected at 24h after initiation of the colonic incubations. $Log_2FC$ is indicated in bold or italic where statistical significance or consistent increase/decrease occurred, respectively. Top 5 species with the largest variations among the treatments are also included in the heat map and indicated with #.

A.

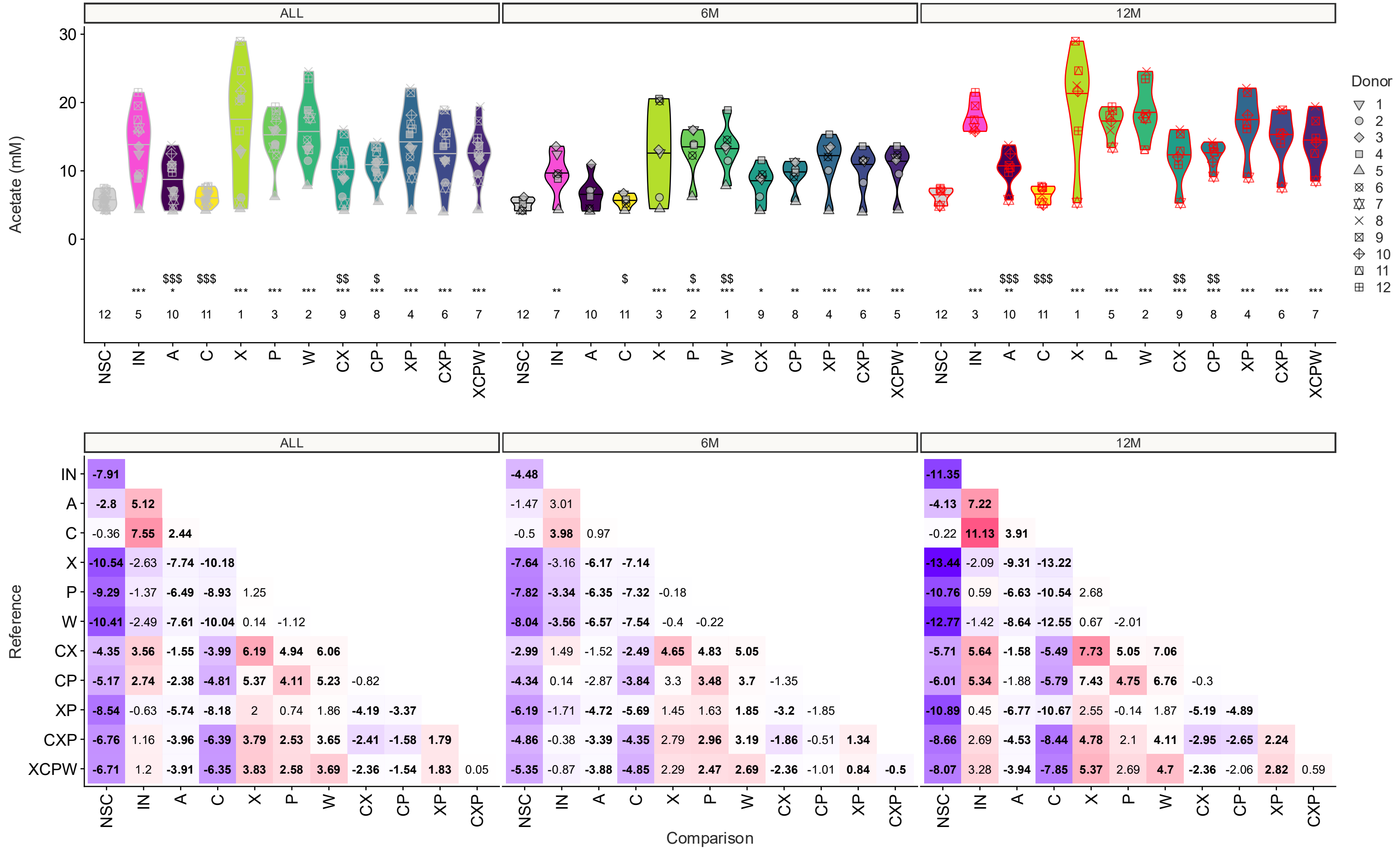

ALL
6M
12M
Acetate (mM)
30
20
10
0
Donor
1
2
3
4
5
6
7
8
9
10
11
12
12 5 10 11 1 3 2 9 8 4 6 7
12 7 10 11 3 2 1 9 8 4 6 5
12 3 10 11 1 5 2 9 8 4 6 7
NSC IN A C X P W CX CP XP CXP XCPW
Reference
Comparison
IN -7.91 | -4.48 | -11.35
A -2.8 5.12 | -1.47 3.01 | -4.13 7.22
C -0.36 7.55 2.44 | -0.5 3.98 0.97 | -0.22 11.13 3.91
X -10.54 -2.63 -7.74 -10.18 | -7.64 -3.16 -6.17 -7.14 | -13.44 -2.09 -9.31 -13.22
P -9.29 -1.37 -6.49 -8.93 1.25 | -7.82 -3.34 -6.35 -7.32 -0.18 | -10.76 0.59 -6.63 -10.54 2.68
W -10.41 -2.49 -7.61 -10.04 0.14 -1.12 | -8.04 -3.56 -6.57 -7.54 -0.4 -0.22 | -12.77 -1.42 -8.64 -12.55 0.67 -2.01
CX -4.35 3.56 -1.55 -3.99 6.19 4.94 6.06 | -2.99 1.49 -1.52 -2.49 4.65 4.83 5.05 | -5.71 5.64 -1.58 -5.49 7.73 5.05 7.06
CP -5.17 2.74 -2.38 -4.81 5.37 4.11 5.23 -0.82 | -4.34 0.14 -2.87 -3.84 3.3 3.48 3.7 -1.35 | -6.01 5.34 -1.88 -5.79 7.43 4.75 6.76 -0.3
XP -8.54 -0.63 -5.74 -8.18 2 0.74 1.86 -4.19 -3.37 | -6.19 -1.71 -4.72 -5.69 1.45 1.63 1.85 -3.2 -1.85 | -10.89 0.45 -6.77 -10.67 2.55 -0.14 1.87 -5.19 -4.89
CXP -6.76 1.16 -3.96 -6.39 3.79 2.53 3.65 -2.41 -1.58 1.79 | -4.86 -0.38 -3.39 -4.35 2.79 2.96 3.19 -1.86 -0.51 1.34 | -8.66 2.69 -4.53 -8.44 4.78 2.1 4.11 -2.95 -2.65 2.24
XCPW -6.71 1.2 -3.91 -6.35 3.83 2.58 3.69 -2.36 -1.54 1.83 0.05 | -5.35 -0.87 -3.88 -4.85 2.29 2.47 2.69 -2.36 -1.01 0.84 -0.5 | -8.07 3.28 -3.94 -7.85 5.37 2.69 4.7 -2.36 -2.06 2.82 0.59
NSC IN A C X P W CX CP XP CXP

B.

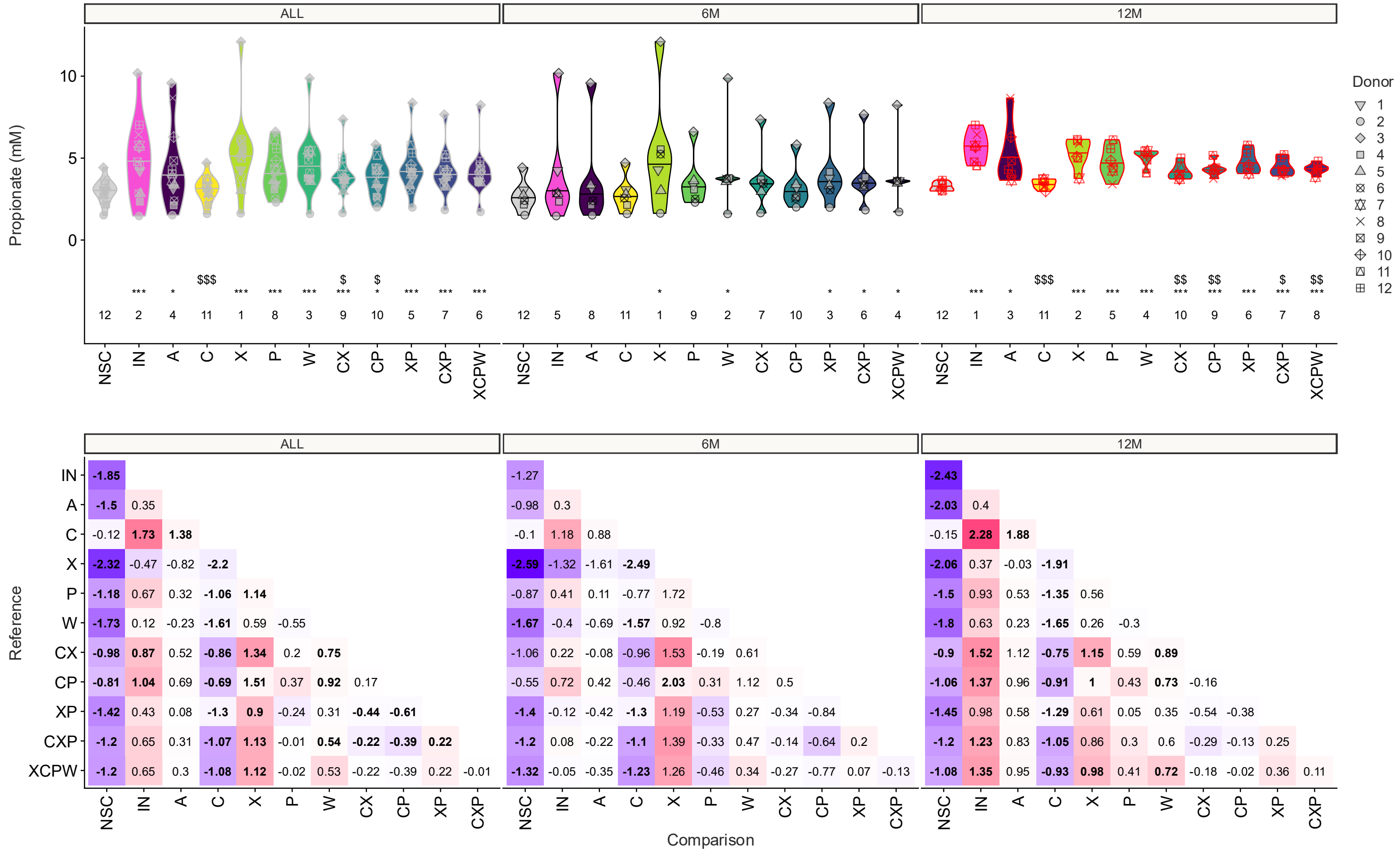
ALL
6M
12M
Propionate (mM)
Donor
NSC
IN
A
C
X
P
W
CX
CP
XP
CXP
XCPW
Reference
Comparison

C.

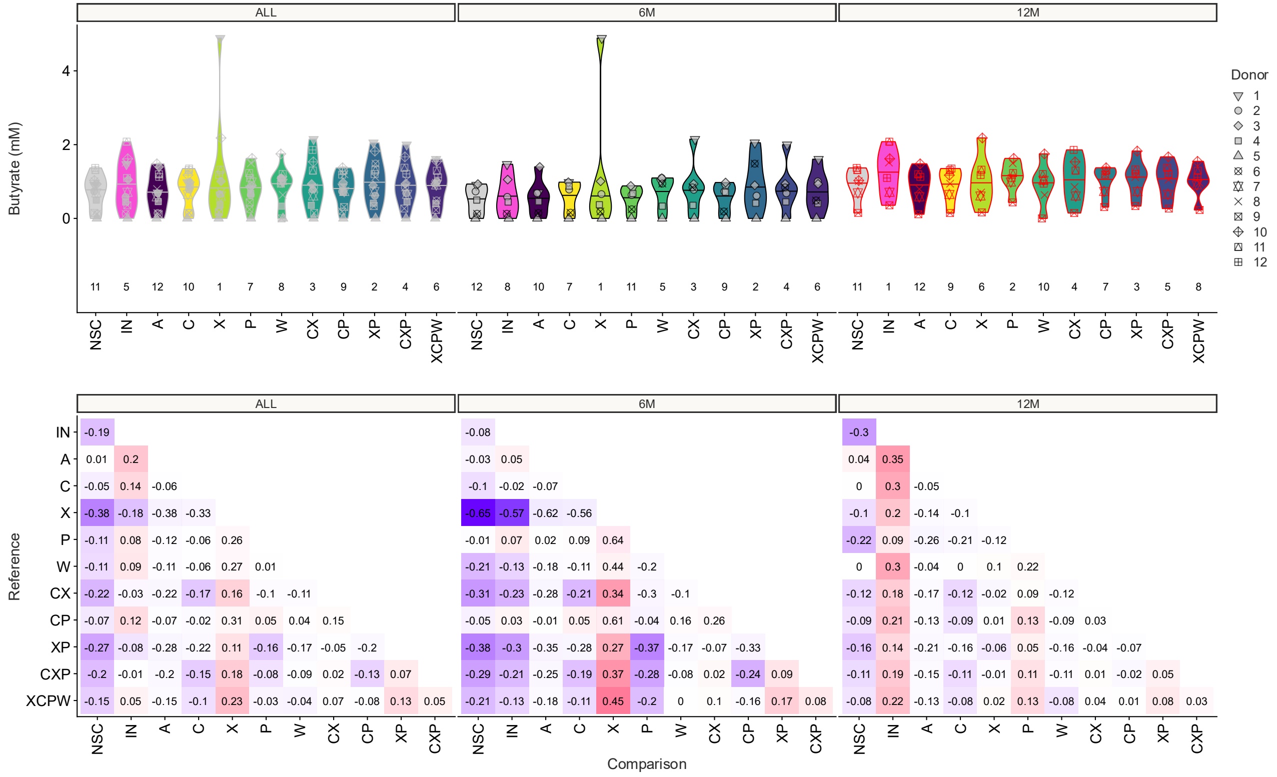


**Supplemental Figure 4. The impact of the tested products on acetate production for 12 infants**

A, Acetate, B. Propionate. C. Butyrate. For each panel, top section shows the impact of the tested products on acetate production for 12 infants (6 months old (n= 6); 12 months old (n = 6)). Statistical differences between NSC and the individual treatments are visualised via * (0.01 < p-adjusted < 0.05), ** (0.001 < p-adjusted < 0.01) or *** (p-adjusted < 0.001). Significant differences between IN and the individual treatments are indicated via $/$$/$$$. The rank of the average values per treatment are indicated at the bottom of the figure. Bottom section listed values in each matrix represent the difference (in mM) between the product on the horizontal axis compared to the one on the vertical axis, with significant differences (p-adjusted < 0.05) being indicated in bold.

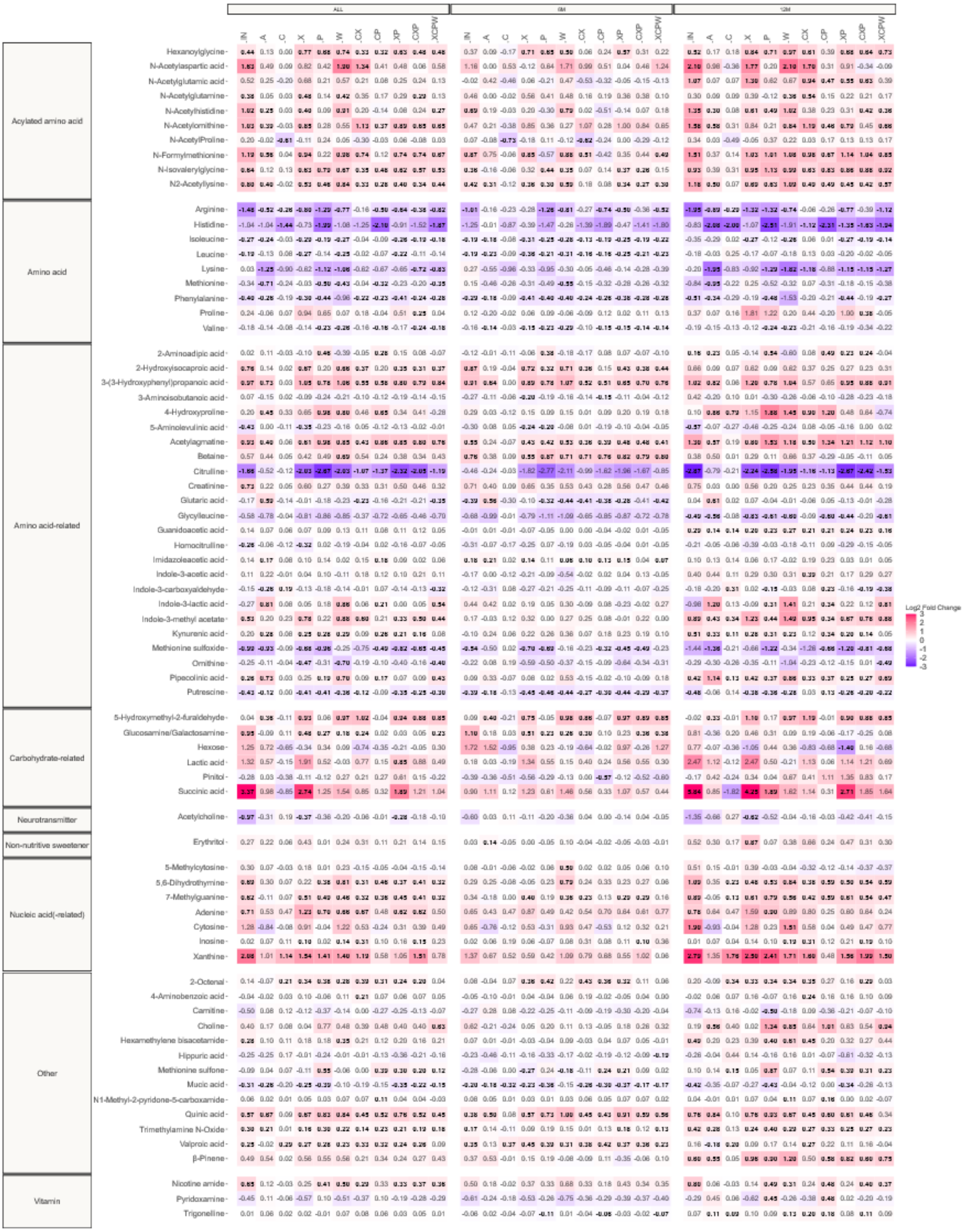


**Supplemental Figure 5.** The impact of the test products compared to a no substrate control (NSC) on metabolites annotated at level 1/2a as quantified via untargeted LC-MS after 24h of incubation, tested via the SIFR® technology for 12 infants (6 months old (n= 6); 12 months old (n = 6)). The metabolites were significantly affected by any of the treatments (p-adjusted < 0.05). Significant differences are indicated by bold of the average log2(abundance treatment/abundance NSC).

A.

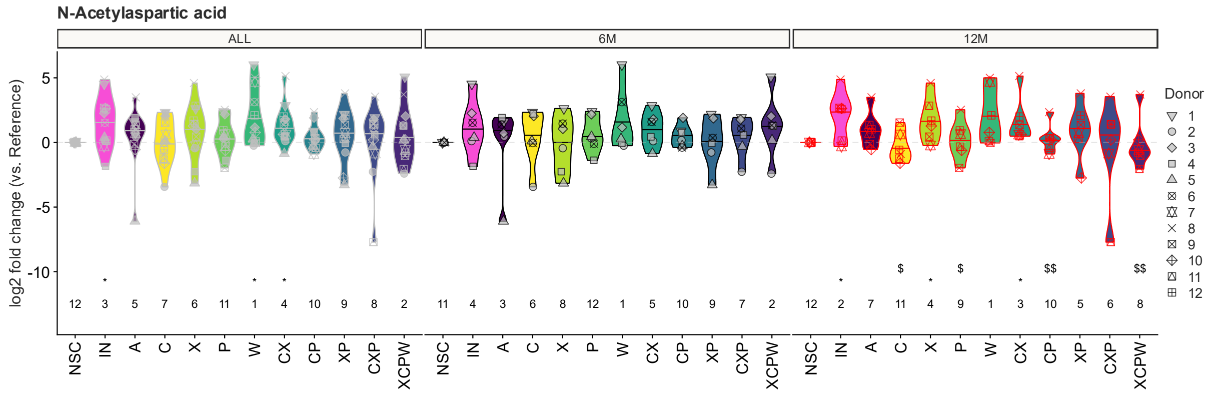


B.

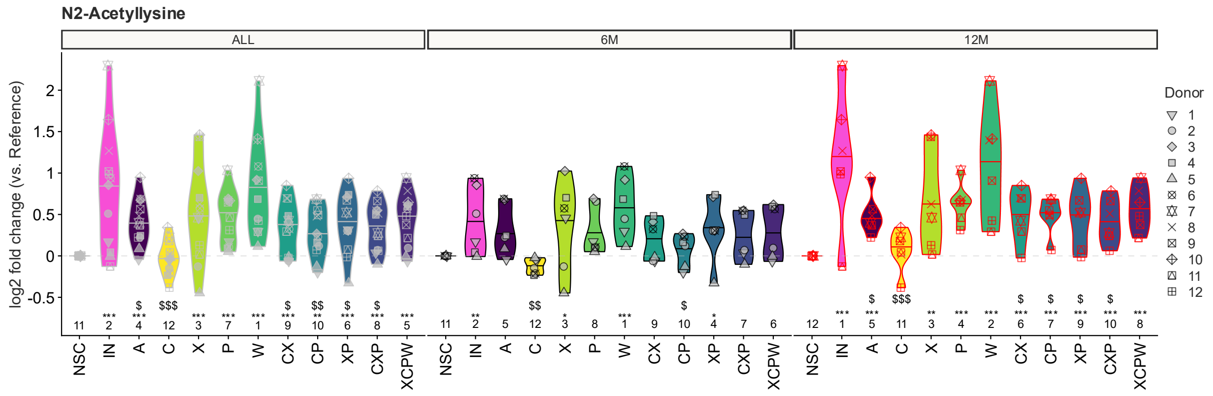


C.

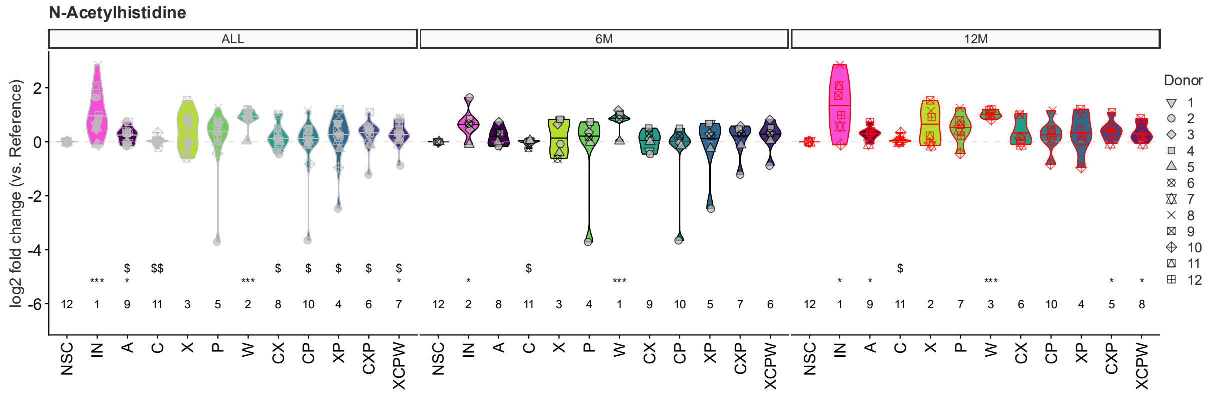


**Supplemental Figure 6.** The impact of the test products on N-acetylaspartic acid (A), N2-acetyllisine (B), and N-acetylhistidine (C) for 12 infants (6 months old (n= 6); 12 months old (n = 6)). Statistical differences between NSC and the individual treatments are visualised via * (0.01 < p-adjusted < 0.05), ** (0.001 < p-adjusted < 0.01) or *** (p-adjusted < 0.001). Significant differences between inulin and the individual treatments are indicated via $/$$/$$$. The rank of the average $Log_2FC$ per treatment are indicated at the bottom of the figure. Ranks comprise here the median of the log-fold changes, not the raw data.

A.

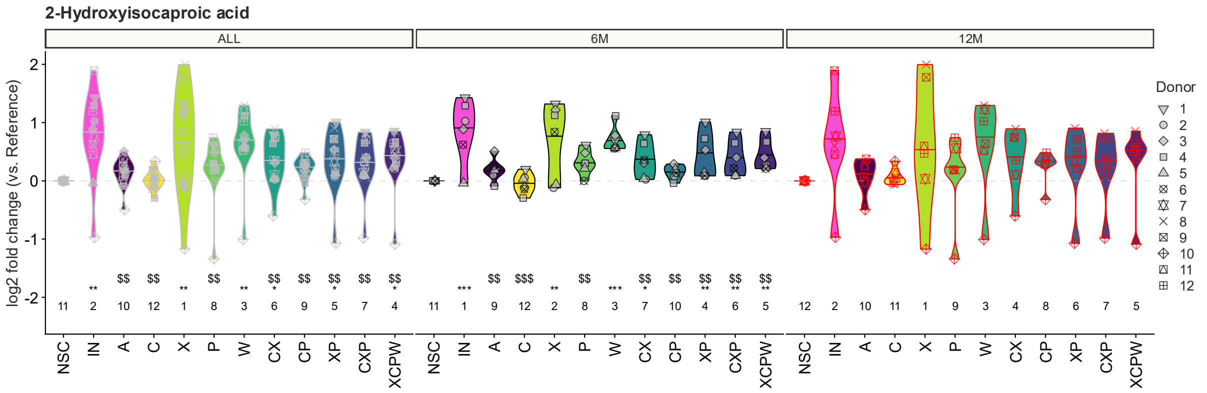
2-Hydroxyisocaproic acid
ALL
6M
12M
log2 fold change (vs. Reference)
NSC IN A C X P W CX CP XP CXP XCPW
Donor
1 2 3 4 5 6 7 8 9 10 11 12

B.

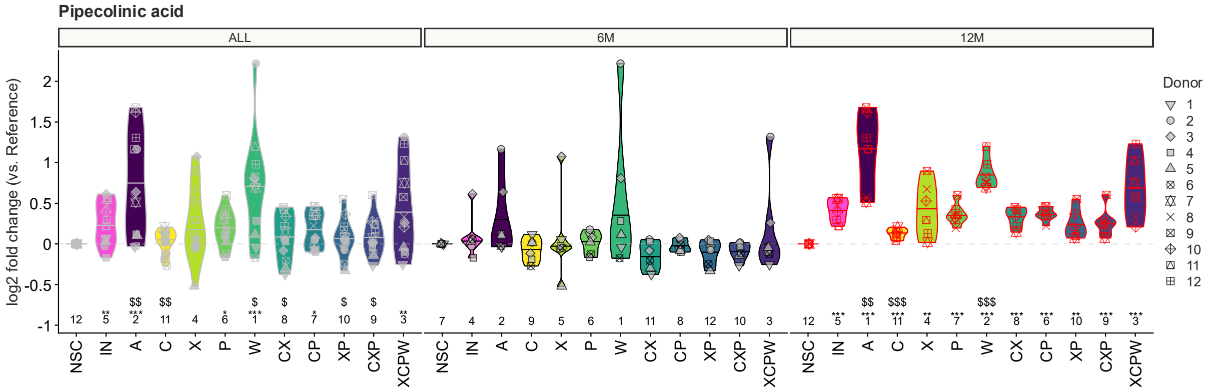
Pipecolinic acid
ALL
6M
12M
log2 fold change (vs. Reference)
NSC IN A C X P W CX CP XP CXP XCPW
Donor
1 2 3 4 5 6 7 8 9 10 11 12

C.

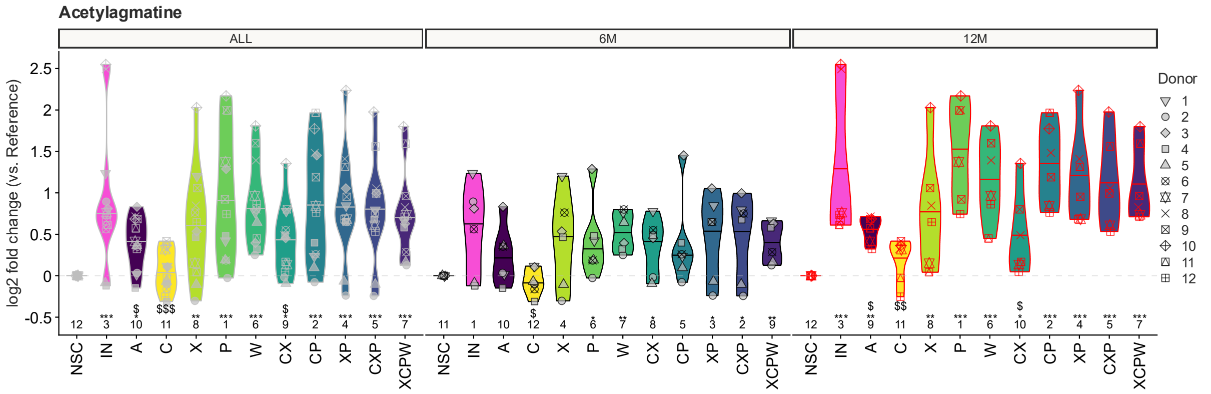
Acetylagmatine
ALL
6M
12M
log2 fold change (vs. Reference)
NSC IN A C X P W CX CP XP CXP XCPW
Donor
1 2 3 4 5 6 7 8 9 10 11 12

D.

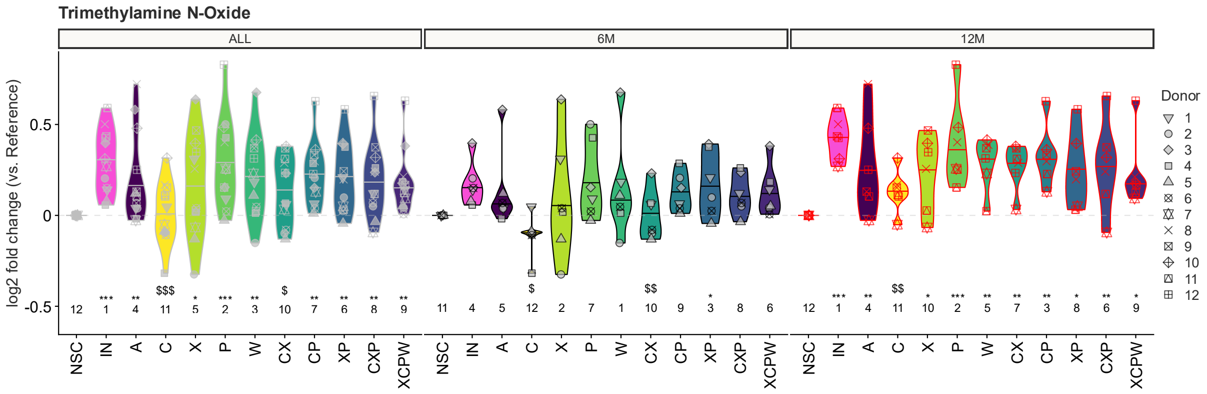


E.

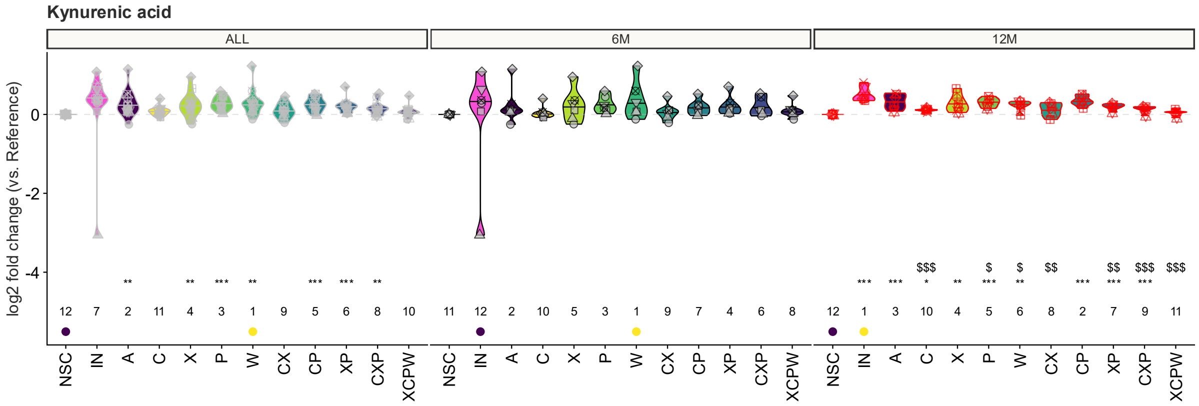


**Supplementary Figure 7.** The impact of the test products on 2-hydroxyisocaproic acid (A), Pipecolinic acid (B), acetylagmantine (C), trimethylamine N-oxide (D) and kynurenic acid (E) for 12 infants (6 months old (n= 6); 12 months old (n = 6)). Statistical differences between NSC and the individual treatments are visualised via * (0.01 < p-adjusted < 0.05), ** (0.001 < p-adjusted < 0.01) or *** (p-adjusted < 0.001). Significant differences between IN and the individual treatments are indicated via $/$$/$$$. The rank of the average $Log_2FC$ per treatment are indicated at the bottom of the figure. Ranks comprise here the median of the log-fold changes, not the raw data.

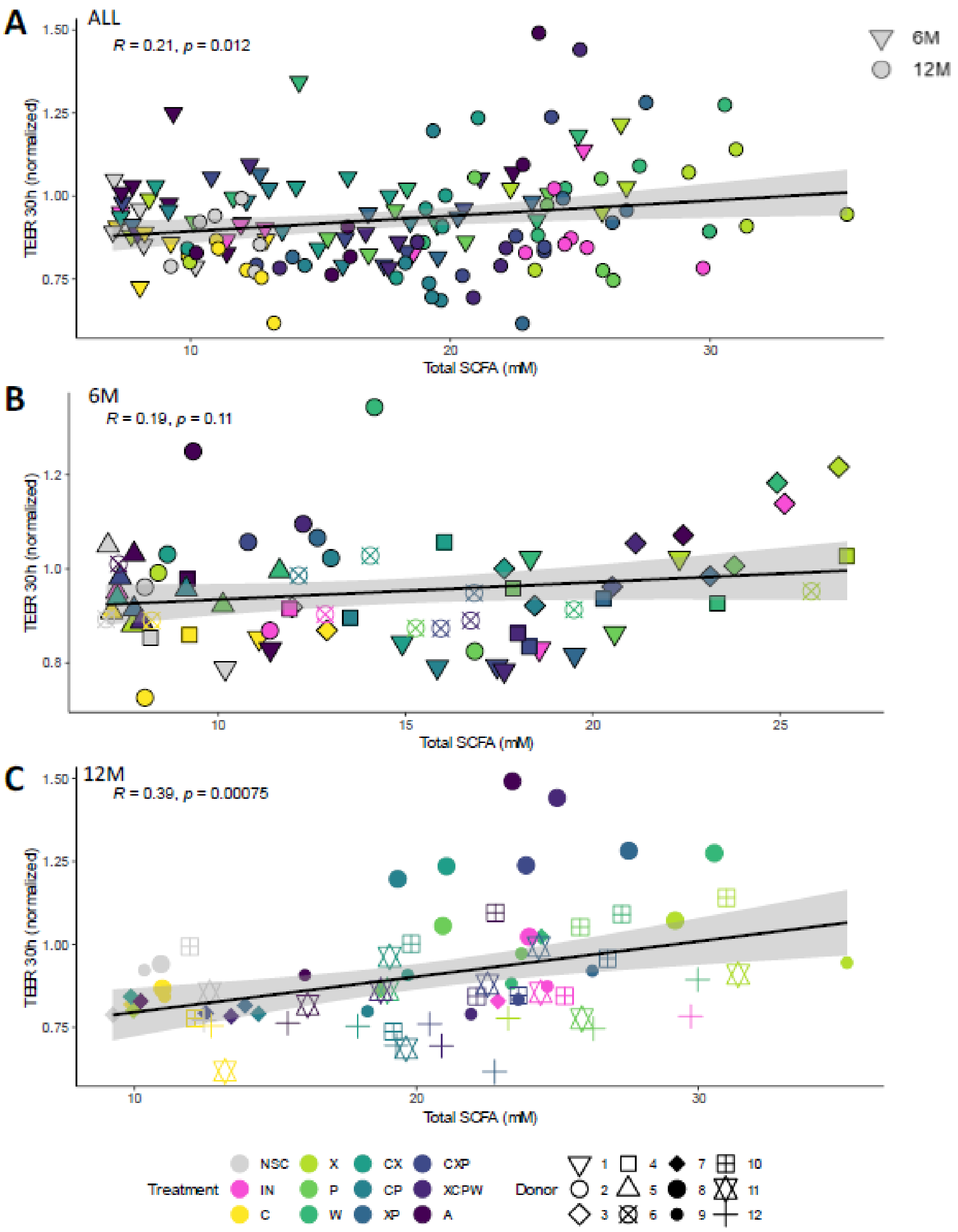


**Supplemental Figure 8**. Correlation analysis between total SCFA and gut barrier integrity as measured by the TEER of the Caco-2 epithelial layer after 30h of treatment (= 24h in absence of LPS and 6h in presence of LPS in the basal compartment containing differentiated THP-1 cells) for (A) all donors (n = 12), (B) 6M infants (n = 6) and (C) 12M infants (n = 6).

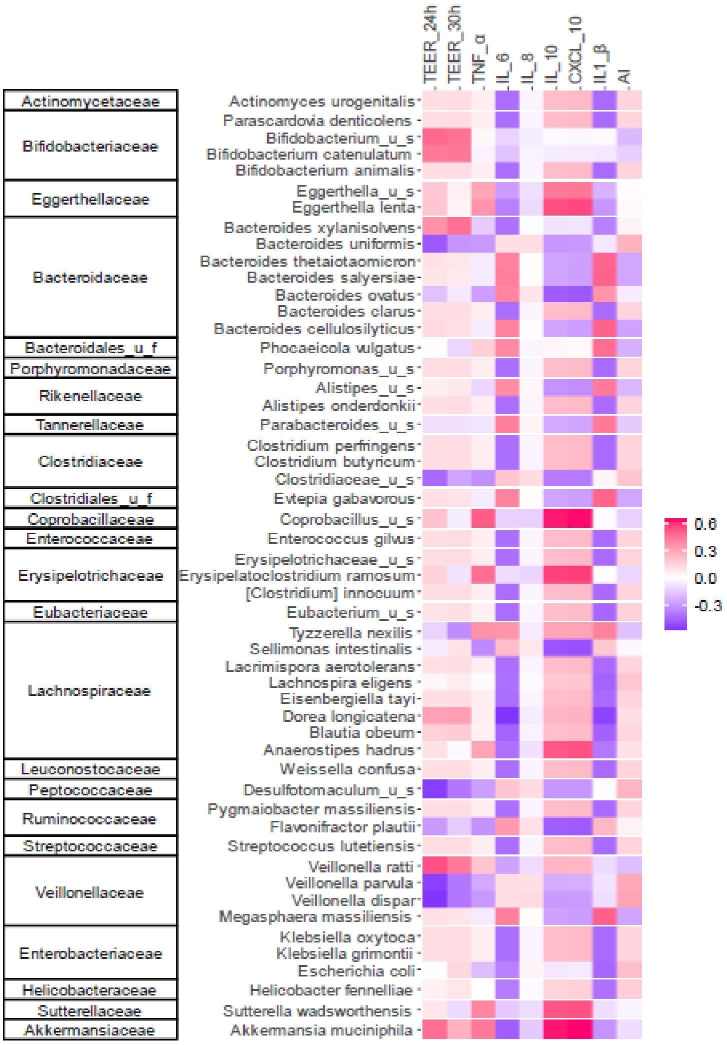


**Supplemental Figure 9**. Regularized Canonical Correlation Analysis (rCCA) to highlight correlations between gut barrier integrity (TEER) and immune markers with microbial composition at species level.

A.

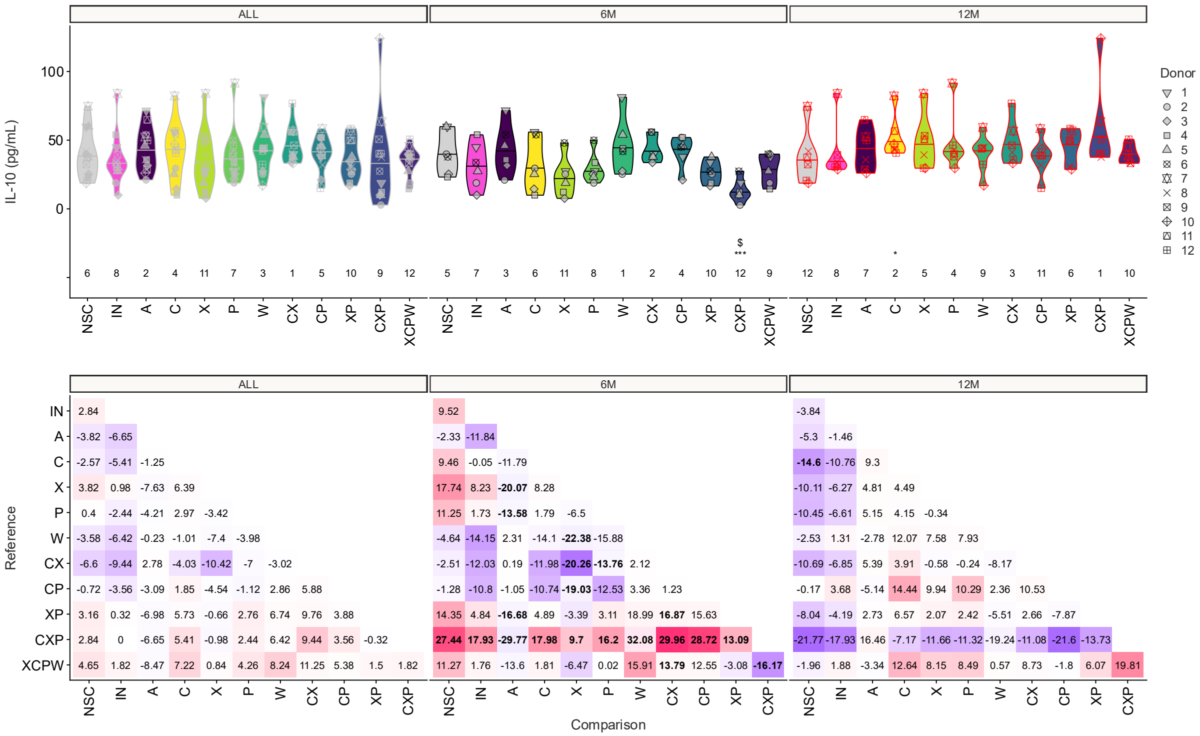
ALL
6M
12M
IL-10 (pg/mL)
Donor
Reference
Comparison
NSC
IN
A
C
X
P
W
CX
CP
XP
CXP
XCPW

**B.**

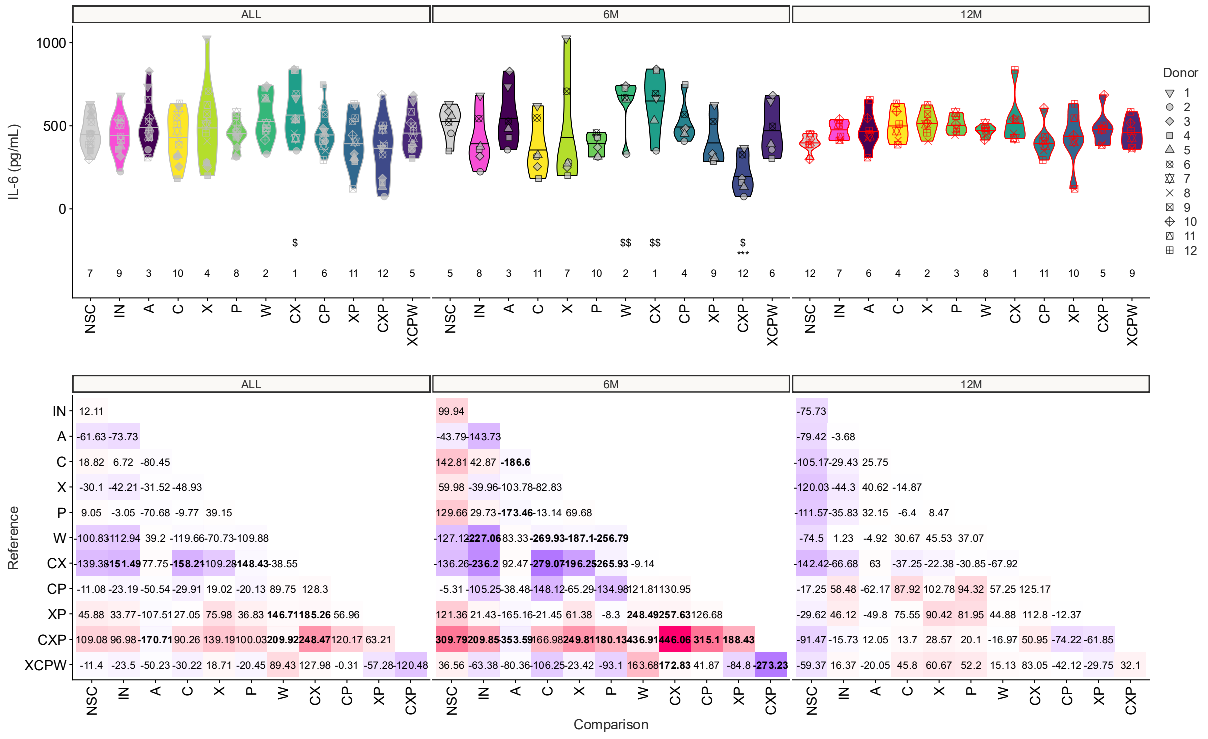


**C.**

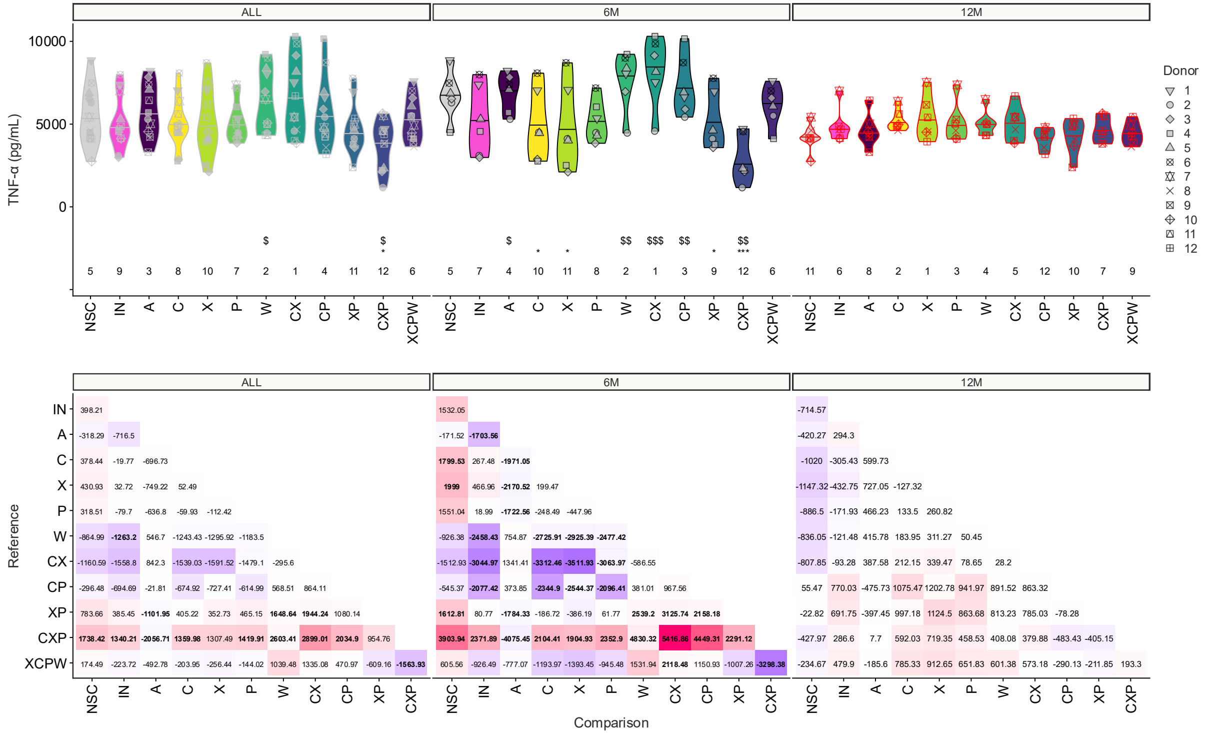
ALL
6M
12M
TNF-α (pg/mL)
10000
5000
0
Donor
1
2
3
4
5
6
7
8
9
10
11
12
NSC
IN
A
C
X
P
W
CX
CP
XP
CXP
XCPW
Reference
Comparison

**D.**

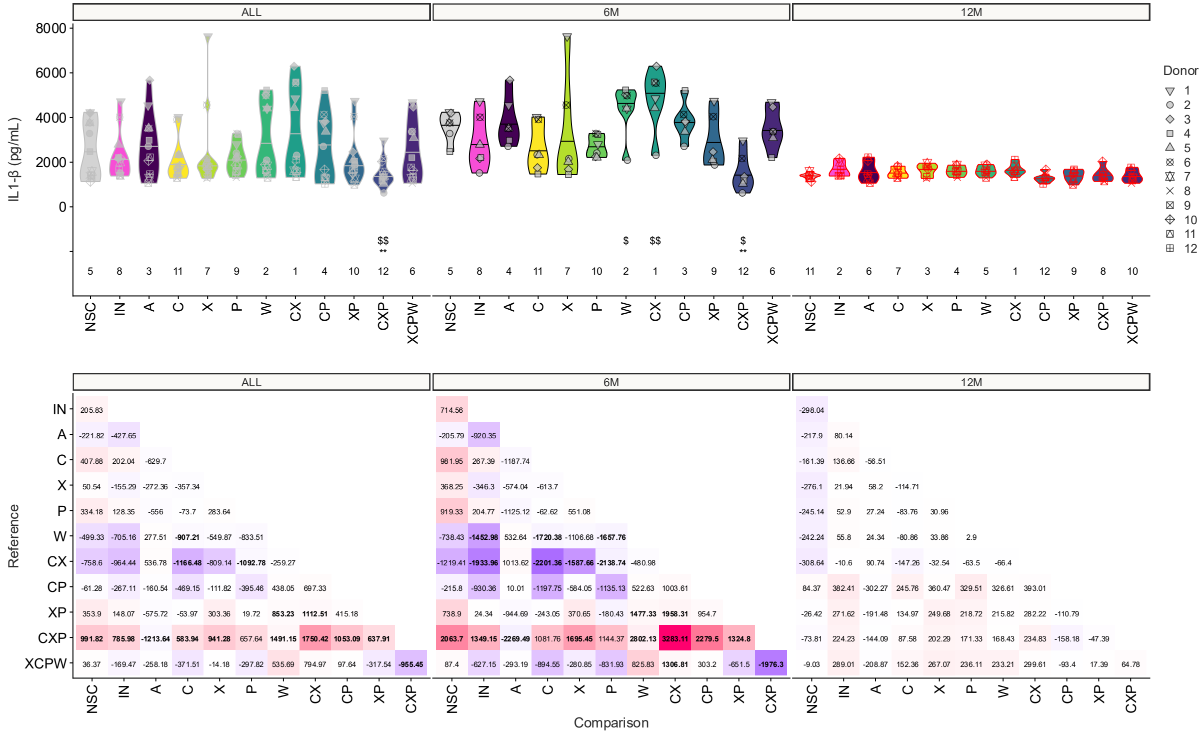

ALL
6M
12M
IL1-β (pg/mL)
8000
6000
4000
2000
0
Donor
NSC
IN
A
C
X
P
W
CX
CP
XP
CXP
XCPW
Reference
Comparison


E.

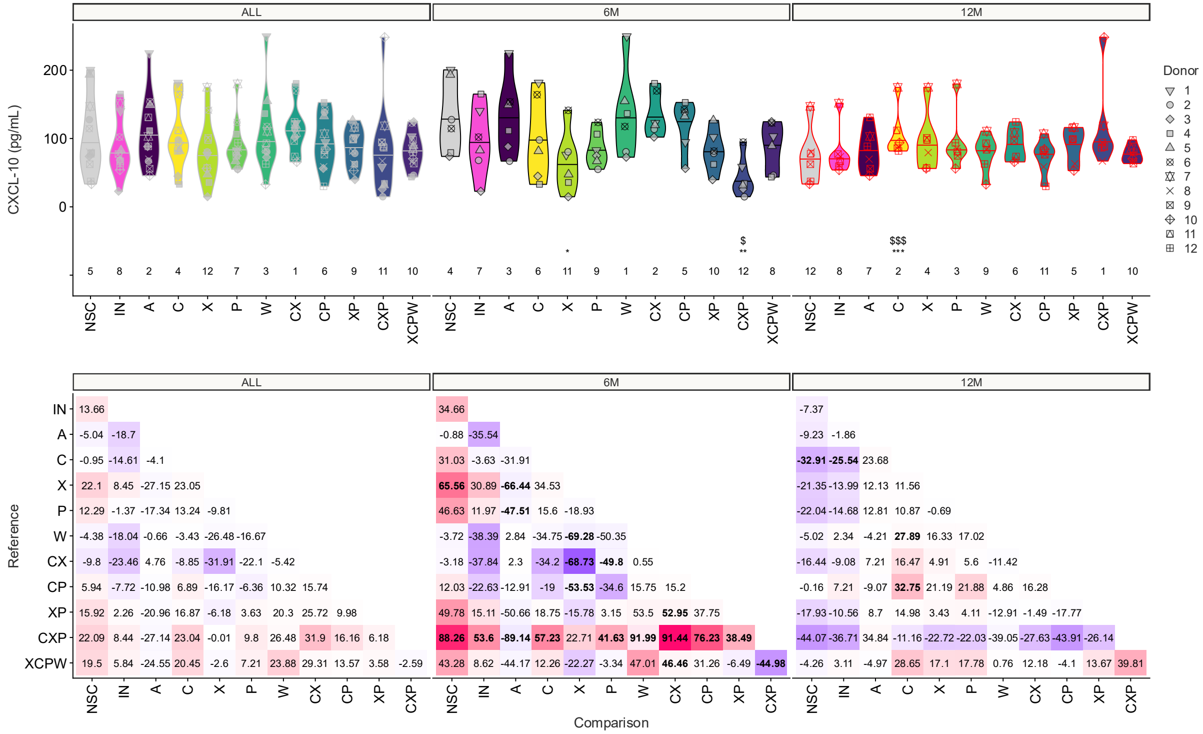


F.

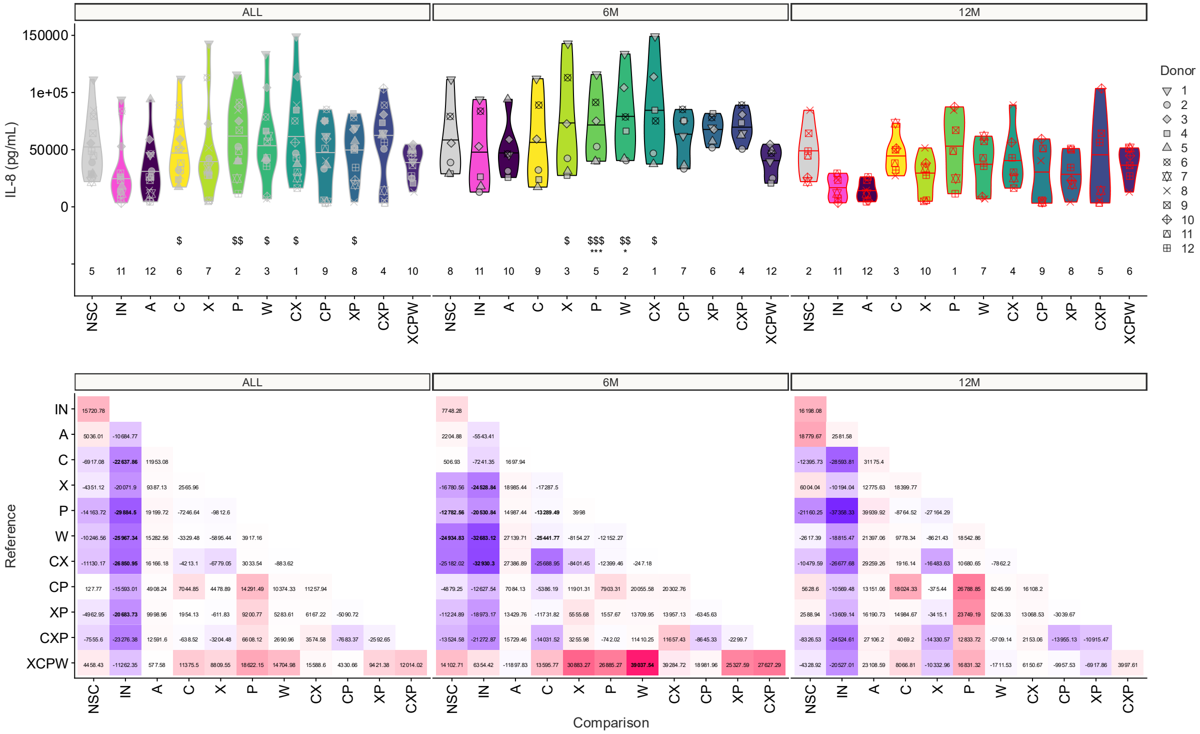


**Supplemental Figure 10.** The impact of the test products on IL-10 (A), IL-6 (B), TNF-α (C), IL1-β (D), CXCL-10 (E), IL-8 (F) for 12 infants (6 months old (n= 6); 12 months old (n = 6)). Statistical differences between NSC and the individual treatments are visualised via * (0.01 < p-adjusted < 0.05), ** (0.001 < p-adjusted < 0.01) or *** (p-adjusted < 0.001). Significant differences between IN and the individual treatments are indicated via $/$$/$$$. The rank of the average value per treatment are indicated at the bottom of the figure. (bottom) Values in each matrix represent the difference (in pg/mL) between the product on the horizontal axis compared to the one on the vertical axis, with significant differences (p-adjusted < 0.05) being indicated in bold.

Supplementary table 2.

The association between pectin intake and microbiota α-diversity in cohort study

| | Simpson | | | | inversimpson | | | | eNS | | | |
|---|---|---|---|---|---|---|---|---|---|---|---|---|
| | Coefficient | 95% CI | | | Coefficient | 95% CI | | | Coefficient | 95% CI | | |
| | | Lower | Upper | P | | Lower | Upper | P | | Lower | Upper | P |
| model1 | 0.027 | 0.017 | 0.036 | <0.001 | 0.553 | 0.255 | 0.851 | <0.001 | 0.75 | 0.346 | 1.154 | <0.001 |
| model2 | 0.01 | 0.001 | 0.02 | 0.039 | 0.334 | 0.005 | 0.663 | 0.047 | 0.414 | -0.029 | 0.857 | 0.068 |
| model3 | 0.017 | 0.004 | 0.03 | 0.013 | 0.577 | 0.126 | 1.027 | 0.012 | 0.726 | 0.119 | 1.333 | 0.019 |
| model4 | 0.007 | 0.003 | 0.011 | 0.001 | 0.191 | 0.052 | 0.331 | 0.008 | 0.261 | 0.074 | 0.449 | 0.007 |

Linear mixed effect models were used for the analysis. Subject number was used as random effect. Model 1 adjusted for age; Model 2 adjusted for age and energy intake; Model 3 adjusted for age, energy and soluble finer intakes; Model 4 each 10% of pectin in total soluble fiber adjusted for age, energy and soluble fiber intakes. Simpson: Simpson index, invsimpson: inverse Simpson index, eNS: expected number of species.
